# Tissue-specific mouse mRNA isoform networks

**DOI:** 10.1101/558361

**Authors:** Gaurav Kandoi, Julie A. Dickerson

**Affiliations:** Bioinformatics and Computational Biology Program, Iowa State University, Ames, IA, USA; Department of Electrical and Computer Engineering, Iowa State University, Ames, IA, USA

## Abstract

Alternative Splicing produces multiple mRNA isoforms of genes which have important diverse roles such as regulation of gene expression, human heritable diseases, and response to environmental stresses. However, little has been done to assign functions at the mRNA isoform level. Functional networks, where the interactions are quantified by their probability of being involved in the same biological process are typically generated at the gene level. We use a diverse array of tissue-specific RNA-seq datasets and sequence information to train random forest models that predict the functional networks. Since there is no mRNA isoform-level gold standard, we use single isoform genes co-annotated to Gene Ontology biological process annotations, Kyoto Encyclopedia of Genes and Genomes pathways, BioCyc pathways and protein-protein interactions as functionally related (positive pair). To generate the non-functional pairs (negative pair), we use the Gene Ontology annotations tagged with “NOT” qualifier. We describe 17 Tissue-spEcific mrNa iSoform functIOnal Networks (TENSION) following a leave-one-tissue-out strategy in addition to an organism level reference functional network for mouse. We validate our predictions by comparing its performance with previous methods, randomized positive and negative class labels, updated Gene Ontology annotations, and by literature evidence. We demonstrate the ability of our networks to reveal tissue-specific functional differences of the isoforms of the same genes.

## Introduction

Recent studies illustrate that genes can have distinct functions with different mRNA isoforms, highlighting the importance of studying mRNA isoforms of a gene [1,2]. This diversity in mRNA isoforms is a result of Alternative Splicing (AS). Many alternatively spliced mRNA isoforms are variably expressed across cell and tissue types [3–10]. AS affects regulation of gene expression, development, human heritable diseases, and response to environmental stresses. This article builds mouse tissue-specific functional networks by integrating heterogeneous expression and sequence datasets at the mRNA isoform level.

In higher organisms such as mouse and human, AS plays a significant role in expanding the variety of protein species [11–14]. As an effect, a gene may produce multiple mRNA isoforms whose protein translations differ in expression, interaction and function [1,5,14,15]. For example, there are more than 75,000 mRNA isoforms encoded by over 20,000 genes in the Mouse genome annotation (GRCm38.p4). The fact that a gene is a mixture of mRNA isoforms makes referencing a gene as being “upregulated” or “downregulated”, uninformative.

Massively parallel sequencing of mRNA isoforms has led to a rapid accumulation of expression and sequence data at the mRNA isoform level. RNA-Seq has provided evidence confirming the production and expression of distinct mRNA isoforms under different conditions [9,16,17]. This has led to the improvement and refinement of genome annotations. Functional networks, at the mRNA isoform level are important for understanding gene function but are largely uninvestigated [18,19].

Traditionally, functional experiments are performed at the gene level. Therefore, there are very few (few hundreds) functional annotations for alternatively spliced mRNA isoforms. The functional data recorded in databases such as Gene Ontology (GO), Kyoto Encyclopedia of Genes and Genomes (KEGG), and UniProt Gene Ontology Annotations (UniProt-GOA) are focused on the canonical mRNA isoform and contain only few hundred annotations describing the functions of alternatively spliced mRNA isoforms. These databases do not store tissue specific information either.

The task of mRNA isoform function prediction is a challenging problem. Some mRNA isoforms are non-functional and introduce noise in the data. Many mRNA isoforms are tissue and condition specific. Since a gene can produce multiple mRNA isoforms [20], the direct transfer of function from the gene to its mRNA isoforms doesn’t work. Gene function prediction methods consider a gene as a single entity. Therefore, these cannot be directly applied to mRNA isoform function prediction because they ignore the distinct functions of alternatively spliced mRNA isoforms. However, important advancements have been made by recent studies towards mRNA isoform level understanding of gene functions [18,19,21–25] such as the prediction of more immune related gene ontology terms for the mRNA isoform ADAM15B than isoform ADAM15A of ADAM15 gene, which is involved in B-cell-mediated immune mechanisms.

One such study developed the human isoform-isoform interactions database (IIIDB) using RNA-Seq datasets, domain-domain interactions and protein-protein interactions (PPIs) [19]. A logistic regression model was built using physical interaction data from the IntAct database [26]. The predicted human isoform-isoform physical interaction network was restricted to the gene pairs already present in IntAct. The problem of mRNA isoform functional network prediction is formulated as a complex multiple instance learning (MIL) problem in [18]. In MIL, a gene is treated as a “bag” of mRNA isoforms (“instances”). A gene pair is formulated as a bag of multiple instance pairs, each of which has different probabilities to be functionally related. The goal of MIL is to identify the specific instance pairs which are functional and maximize the difference between them and the instance pairs of non-functionally related bags. A Bayesian network based MIL algorithm was developed by [18] to predict a mouse mRNA isoform level functional network using RNA-Seq datasets, exon array, pseudo-amino acid composition and isoform-docking data.

The studies [18,19] above introduce bias in the training and testing dataset by using random mRNA isoform pairs as non-functional pairs (negative pairs) and do not consider the tissue-specific mRNA isoform functions. Our work is fundamentally different and improves upon the studies [18,19] above both in terms of research content and computational approaches. First, we formulate the problem of mRNA isoform functional network prediction as a simple supervised learning task. Second, our goal is to develop tissue-specific functional networks for mouse. Lastly, like previous methods, we do not introduce bias by assuming that functionally unrelated (negative pair) mRNA isoform pairs can be selected based on the cellular localization [19] or at random [18], which is crucial to the selection of training data in a machine learning system.

We have developed 17 tissue-specific mRNA isoform level functional networks in addition to an organism level reference functional network for mouse. Using the leave-one-tissue-out strategy with a diverse array of tissue-specific RNA-Seq datasets and sequence information, we trained a random forest model to predict the functional networks. Because there is no mRNA isoform-level gold standard for testing, we have used the single mRNA isoform genes co-annotated to GO biological process, KEGG pathways, BioCyc pathways and PPIs as functionally related (positive pair). The non-functional pairs (negative pairs) were generated by using the GO annotations tagged with “NOT” qualifier. We have validated our predictions by comparing its performance with previous methods, datasets with randomized positive and negative class labels, updated GO annotations and literature evidence.

## Methods

### mRNA isoform level data processing

This study considers mRNA isoforms annotated in the NCBI *Mus musculus* genome assembly (GRCm38.p4) for which both mRNA and protein sequences are available. All protein (and corresponding mRNA) sequences smaller than 30 amino acids and those containing non-standard characters are not considered. This resulted in a filtered set of 75,826 mRNA isoforms from 21,813 genes.

To comprehensively characterize mRNA isoform pairs, we have processed 359 RNA-Seq samples from 17 tissues and calculated protein and mRNA sequence properties as described below. Such heterogeneous features have been shown to be useful for predicting several biological properties [18,27,28]. All calculations and analyses were performed on the Extreme Science and Engineering Discovery Environment (XSEDE) Comet cluster [29].

The mRNA and protein level features are summarized in Table 1 and an overview of the workflow is presented in Fig 1.

**Table 1.**
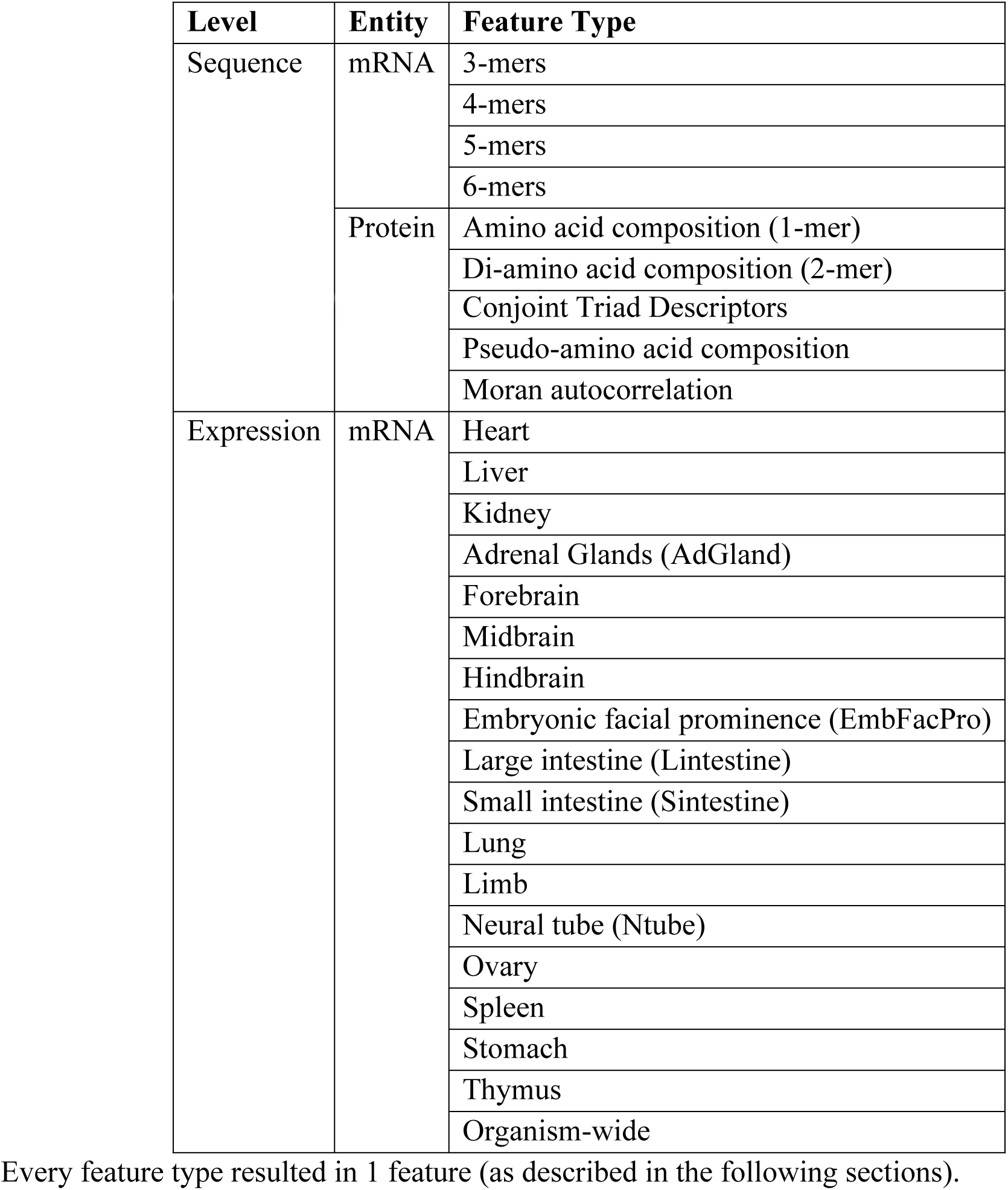
A list of all mRNA and protein level feature types used in this study.

**Fig 1.**
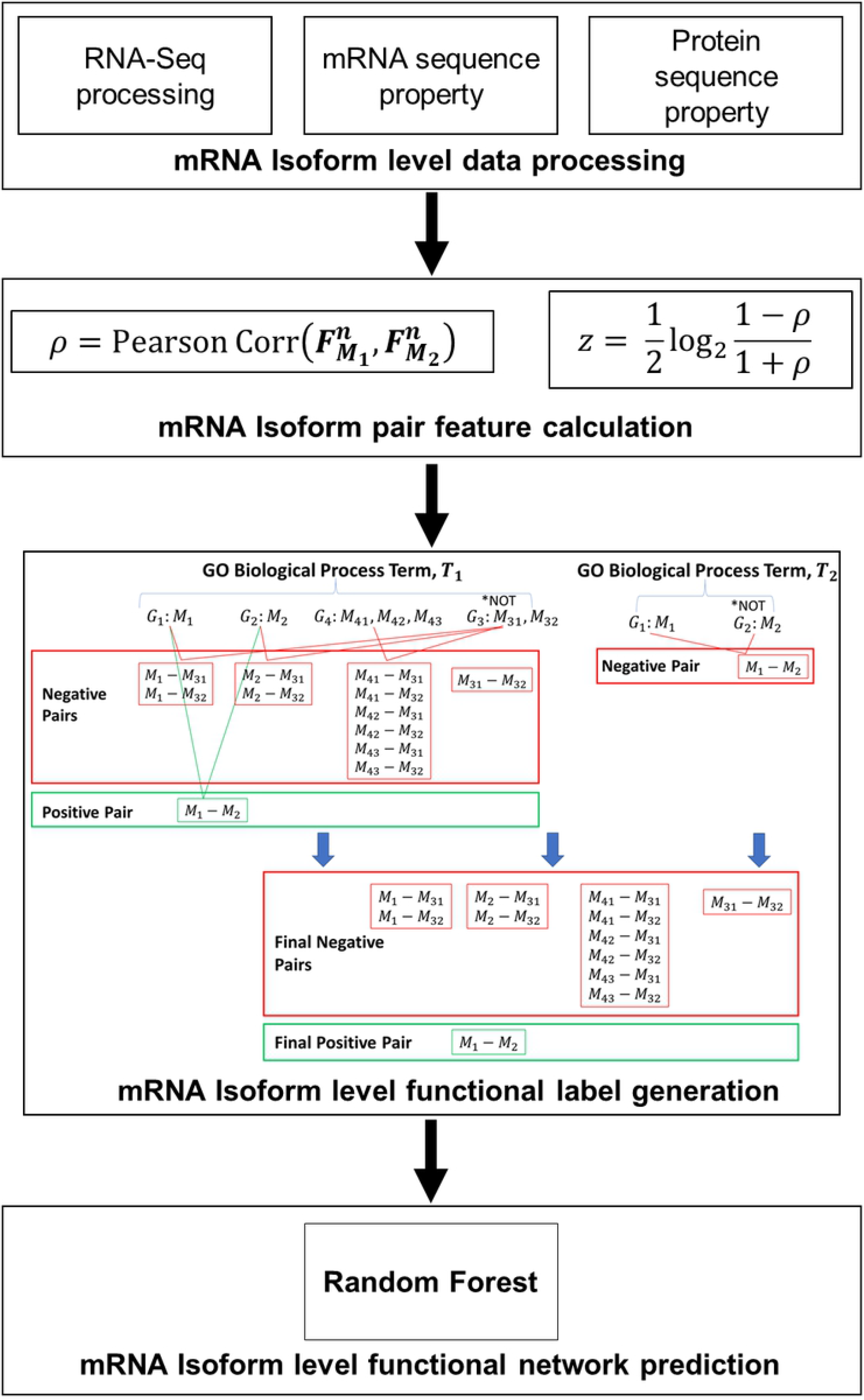
Overview of our workflow. A brief overview of TENSION is provided. We also illustrate the process of generating the mRNA isoform level labels using two dummy gene ontology biological process terms, T_1_ and T_2_. Functional mRNA isoform pairs (positive pairs) are shown in green and non-functional pairs (negative pairs) are shown in red.

#### Preprocessing of RNA-seq datasets

To capture tissue specific functions, RNA-Seq datasets from multiple tissues are processed to extract the expression values. Starting with the ENCODE mouse RNA-Seq datasets, the following filtering criteria are used to select the datasets: 1) Read length ≥ 50; 2) Mapping percent ≥ 70%; and 3) No error or audit warning flags were generated. For the tissue specific networks, only those tissues with at least 10 samples were used. Based on these filters we retained 359 RNA-Seq samples from around 20 tissues, 17 of which have at least 10 samples (Table S1).

The mouse genome build GRCm38.p4 from NCBI was used to align the RNA-Seq datasets using STAR (version 2.5.3a) [30]. Then, the relative abundance of the mRNA isoforms as fragments per kilobase of exon per million fragments mapped (FPKM) is calculated using StringTie (version 1.3.3b) [31]. The GFF3 annotation file corresponding to the GRCm38.p4 build was also used during the alignment and quantification.

#### mRNA sequence composition

mRNA sequences can be represented as the frequencies of *k* neighboring nucleic acids, jointly referred to as *k-mers*. For an mRNA sequence there are **4**^*k*^ possible *k-mers* in a *k-mer group*, while there are **20**^*k*^ possible *k-mers* for protein sequences. For a sequence of length *l*,

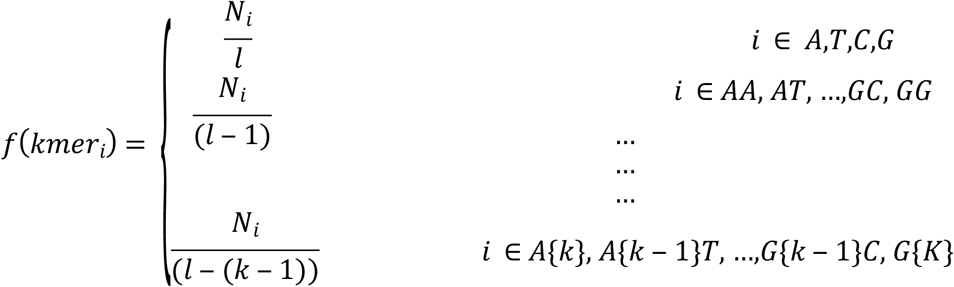

where, *f*(*kmer_i_*) is the frequency of the *ith k-mer* and *N_i_* is the count of the *ith k-mer*. We compute the *k-mer* composition for *k = 3 to 6* for all mRNA isoform sequences using the rDNAse library in R [32,33].

#### Protein Sequence Properties

Each protein sequence can be characterized in multiple ways by exploiting its sequence and order composition. Like the mRNA sequence k-mer composition described above, we compute the k-mer compositions for *k = 1 and 2* for all protein sequences. We also compute the conjoint triad descriptors [34] for all protein sequences. For this, the standard 20 amino acids are grouped into 7 classes according to the volume of the side chains and their dipoles. Then, the *k-mer* composition is calculated at *k = 3* for this newly represented protein sequence. To take the sequential information of the amino acids in a protein sequence into account, we also compute the pseudo-amino acid composition [35] and Moran autocorrelation [36] for all protein sequences. All protein sequence properties were computed using the protr library in R [32,37].

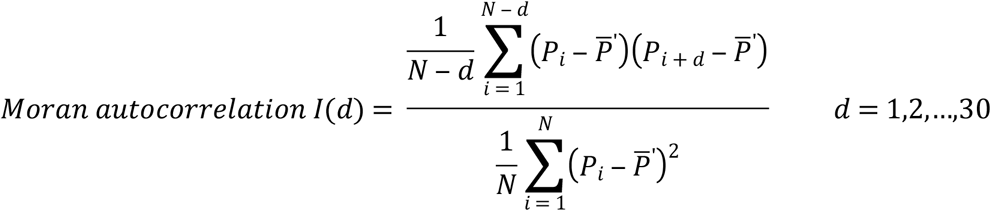

where, *d* is called the lag of the autocorrelation; *P_i_* and *P_i + d_* are the properties of the amino acid *i* and *i* + *d*; 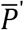 is the considered property *P* along the sequence, i.e.,

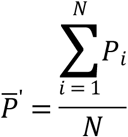

### mRNA isoform level feature calculation

The goal is to accurately predict a functional network which represents the probability of two mRNA isoforms belonging to the same GO biological process or pathway. Lower edge weights correlate with mRNA isoform pairs’ involvement in the same GO biological process or pathway. The weighted functional network is modeled as a graph *G* = (*V,E*), where the set *V* represents the mRNA isoforms (nodes) and the set *E* represents the mRNA isoform pairs (edges). For an mRNA isoform pair (*E_ij_*) in the functional network, the class label (*L_ij_*) is assigned as following:

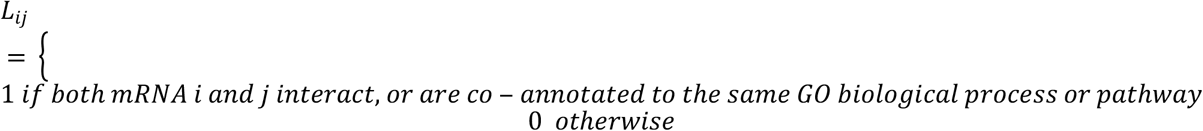

Many mRNA isoforms have zero FPKM values. The FPKM values were corrected by performing log-transformation, i.e. log_2_ (*FPKM* + 1). This alleviates the problem where the log of zero FPKM value is not defined, which is not an acceptable input for machine learning methods.

For all mRNA isoform pairs, Fisher’s z-transformed Pearson correlation scores are calculated and used as input features for machine learning.

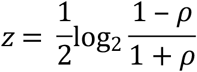

Pearson correlation coefficient of 1 and −1 leads to z-score of -∞ and ∞ respectively, so these z-scores are replaced with an extreme value of −100 and 100 respectively. In cases where the Pearson correlation coefficient is not defined, we set the z-score to 0.

For every mRNA isoform pair, we calculate one z-score using the samples from one tissue and use this as one feature. For instance, one z-score for heart, one z-score for liver, one z-score for lungs and so on for all 17 tissues. Additionally, one z-score is also calculated using all 359 RNA-Seq samples. This resulted in 18 features, one for each of the 17 tissues and one using all RNA-Seq samples. Similarly, for every mRNA isoform pair, we calculate one z-score for each of *k* = 3, 4, 5, and 6 for mRNA isoform sequences, *k* = 1 and 2 for protein sequences, conjoint-triad descriptors, pseudo-amino acid composition and Moran autocorrelation. This led to 9 further features resulting in a total of 27 features.

### mRNA isoform level functional labels

The mRNA isoform level functional labels are created by combining the information from GO biological process annotations (downloaded on 23 October 2017), KEGG pathways (downloaded on 25 September 2017), BioCyc pathways (downloaded on 25 September 2017) and PPIs (downloaded on 25 September 2017). We remove all GO biological process annotations with the evidence codes: Inferred from Electronic Annotation (IEA), Non-traceable Author Statement (NAS) and No biological Data available (ND). We utilize the GO hierarchy (gene ontology downloaded on 25 October 2017) and propagate all annotations by following the “true path rule”, which means that all genes/proteins annotated to a GO term *T* will also be annotated to all ancestor terms of *T.*

The PPIs were integrated from IntAct [26], Biological General Repository for Interaction Datasets (BioGRID) [38], Agile Protein Interactomes DataServer (APID) [39], Integrated Interactions Database (IID) [40] and Mentha [41]. For APID [39], we include interactions with at least 2 experimental evidences (level 2 dataset). For IID [40], we remove all interactions for which there is only orthologous evidence. For Mentha [41], we remove interactions with a score less than 0.2. Finally, we consider PPIs only if both interactors are from mouse.

After propagation, we remove the GO biological process terms which are too broad (more than 1000 genes annotated) or too specific (less than 10 genes annotated). A gene is assumed to be functional if it is annotated to a GO biological process or a pathway. Two genes are assumed to be functionally related if both are co-annotated to the same GO biological process or pathway. The information in GO, KEGG, BioCyc and PPI databases usually focus on the canonical form of a gene/protein and doesn’t distinguish between the mRNA isoforms resulting from AS. So, we construct mRNA isoform level functional labels by utilizing the information from single mRNA producing genes and gene annotations tagged with a “NOT” qualifier. A summary of the mRNA isoform level functional label generation is illustrated in Fig 1.

In our functional networks, if a gene *G*_1_ produces only a single mRNA *M*_1_, then *M*_1_ is assumed to perform the functions of *G*_1_ and is considered functional. Similarly, if two genes *G*_1_ and *G*_2_, both of which produce single mRNAs, *M*_1_ and *M*_2_ respectively, are co-annotated to the same GO biological process or pathway, the pair (edge) *M*_1_ – *M*_2_ is assumed to be functionally related (positive pair). Additionally, if *G*_1_ and *G*_2_ are involved in a PPI, the pair (edge) *M*_1_ – *M*_2_ is also assumed to be functionally related (positive pair).

We utilize a more robust way of defining functionally unrelated (negative pair) mRNA isoform pairs by using the GO biological process annotations tagged with “NOT” qualifier. A gene/protein tagged with “NOT” qualifier means that it is not involved in the respective GO biological process and hence can be considered non-functional (negative) for this GO biological process. All such annotations are propagated by the inverse of “true path rule”, which means that if a gene/protein is explicitly ‘NOT’ annotated to a GO term T, it will also be ‘NOT’ annotated to all the children of *T*. Consider a GO biological process term *T*_1_ annotated with genes *G*_1_ *G*_2_,*G*_3_ and *G*_4_ which produce mRNA isoforms *M*_1_,*M*_2_, *M*_31_, *M*_32_, *M*_41_, *M*_42_, and *M_43_.* Of these genes, if *G*_3_ is tagged with a ‘NOT’ qualifier (Fig 1), all pairs of *M*_31_ and *M*_32_ with *M*_1_ *M*_2_, *M*_41_, *M*_42_, and *M*_43_ are assumed to be functionally unrelated (negative pair). It should be noted that currently there are only few hundred such annotations.

Genes can be annotated to multiple GO biological process terms. In Fig 1, single mRNA isoform producing genes *G*_1_ and *G*_2_ are annotated to GO biological process terms *T*_1_ and *T*_2_. However, the gene *G*_2_ is tagged with a “NOT” qualifier for term *T*_2_. Consequently, the mRNA isoform pair *M*_1_ – *M*_2_ is functionally related for term *T*_1_ but functionally unrelated for term *T*_2_. In cases where an mRNA isoform pair (*M*_1_ – *M*_2_) is found to be both functionally related (positive pair) for one term (*T*_1_) but functionally unrelated (negative pair) for another term (*T*_2_), we consider the mRNA isoform pair (*M*_1_ – *M*_2_) as functionally related (positive pair) because *M*_1_(*G*_1_) and *M*_2_(*G*_2_) are involved in at least one common GO biological process.

### Predicting functional networks

#### Generating training and testing datasets

There are approximately 2.9 billion possible mRNA isoform pairs resulting from the 75,826 annotated mRNA isoforms. Using the method described above (see methods section ‘mRNA isoform level functional labels’), we labelled 2,083,679 mRNA isoform pairs as functional pairs (positive) and 818,071 mRNA isoform pairs as nonfunctional pairs (negative). All the remaining mRNA isoform pairs are considered to be ‘unknown’, i.e. neither functional nor non-functional pairs. The mRNA isoform pairs in the functional and non-functional groups are mutually exclusive, i.e. an mRNA isoform pair can be either functional or non-functional, but not both.

We generate two types of datasets: training and testing. The training and testing datasets are mutually exclusive, i.e. an mRNA isoform pair can be either in a training or testing dataset, but not both. The training dataset contains randomly selected 640,000 functional and 640,000 nonfunctional mRNA isoform pairs. The testing dataset contains randomly selected 160,000 functional and 160,000 non-functional mRNA isoform pairs not included in the training dataset. The functional pairs in the original testing dataset are made up of only single mRNA isoform genes. The non-functional pairs are however not restricted to single mRNA isoform genes. All datasets are balanced.

#### Random forest model for the functional networks

We formulate the task of mRNA isoform functional network prediction as a simple supervised learning problem. In supervised learning, a model capable of distinguishing a pre-defined set of ‘positives’ (functional mRNA isoform pairs in our case) from a set of ‘negatives’ (non-functional mRNA isoform pairs in our case) is built using a set of features derived from potential predictors of the property under consideration (mRNA isoform pair function in our case).

Using all 27 features for our training dataset, we train a Scikit-learn [42] Random Forest [43] model to predict the mRNA isoform functional network. Then we evaluate the performance of the random forest model by making predictions on the testing dataset. Commonly used performance evaluation metrics such as Accuracy, Area Under the Receiver Operating Characteristics Curve (AUROC), Area Under the Precision-Recall Curve (AUPRC), Precision, Recall, F1 Score, and Matthews Correlation Coefficient (MCC) are calculated using the predictions for testing dataset to assess the performance of the random forest model. The predictions are only evaluated when all 27 features are used for predictions. Finally, we use the random forest models to make predictions on all 2.9 billion possible mRNA isoform pairs.

#### Building tissue-specific mRNA isoform networks

To build the tissue-specific mRNA isoform networks, we utilize the leave-one-tissue-out strategy. First, using all 27 features, we train an organism-level mRNA isoform functional network prediction random forest model. Then, we generate 17 tissue-specific mRNA isoform functional network prediction random forest models by removing the tissue specific RNA-Seq features, one tissue at a time. The mRNA isoform pairs for which the prediction is unaffected after leave-one-tissue-out are referred to as “reference pairs”. The two tissue-specific cases are: 1) mRNA isoform pairs which are predicted to be functional in only one tissue (tissue specific functional mRNA isoform pairs), and 2) mRNA isoform pairs which are predicted to be non-functional in only one tissue (tissue specific non-functional mRNA isoform pairs). These are also summarized in Fig 2.

**Fig 2.**
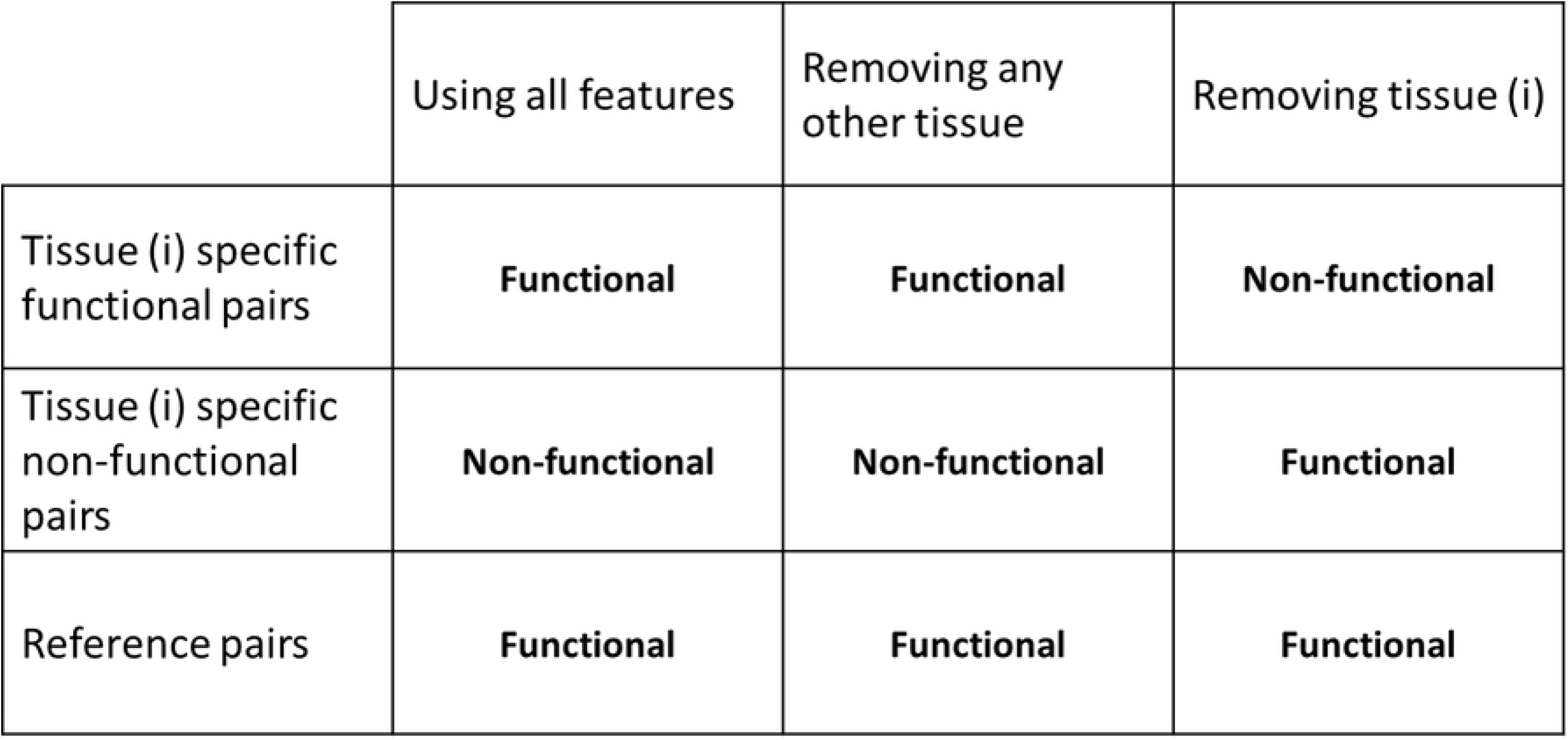
Defining tissue specific functional and non-functional mRNA isoform pairs. Here we illustrate the process of classifying the mRNA isoforms as tissue specific functional, tissue specific non-functional or organism wide reference pairs. If the prediction is functional (positive) when using all 27 features but changes to non-functional (negative) after removing the tissue derived RNA-Seq feature, we assume such mRNA isoform pairs as tissue-specific functional pairs. Contrary to tissue-specific functional pairs, if the prediction changes from non-functional (negative) to functional (positive) after removing the tissue derived RNA-Seq feature, we assume such pairs as tissue-specific non-functional pairs. For the reference pairs, the prediction is constant after removing any tissue derived RNA-Seq feature.

If the prediction for an mRNA isoform pair changes from functional (positive) to non-functional (negative) after removing a tissue derived RNA-Seq feature, we consider such mRNA isoform pairs as tissue specific functional pairs. Similarly, if the prediction for an mRNA isoform pair changes from non-functional (negative) to functional (positive) after removing a tissue derived RNA-Seq feature, we consider such mRNA isoform pairs as tissue specific non-functional pairs. For instance, consider the case of heart specific mRNA isoform functional network prediction. We train two random forest models, 1) using all 27 features and, 2) after removing the heart derived RNA-Seq feature. Then, the heart specific functional mRNA isoform pairs are those which are predicted as functional by the first model but non-functional by the second model and vice-versa for the non-functional mRNA isoform pairs.

#### From mRNA isoform networks to gene networks

We collapse the tissue-specific mRNA isoform networks to gene networks as illustrated in Fig 3. All mRNA isoform nodes of the same gene are merged into a single gene node. All direct edges from the mRNA isoforms of the same gene are transferred to the single gene node. This resulted in 17 gene level tissues networks in addition to the 17 tissue-specific mRNA isoform networks.

**Fig 3.**
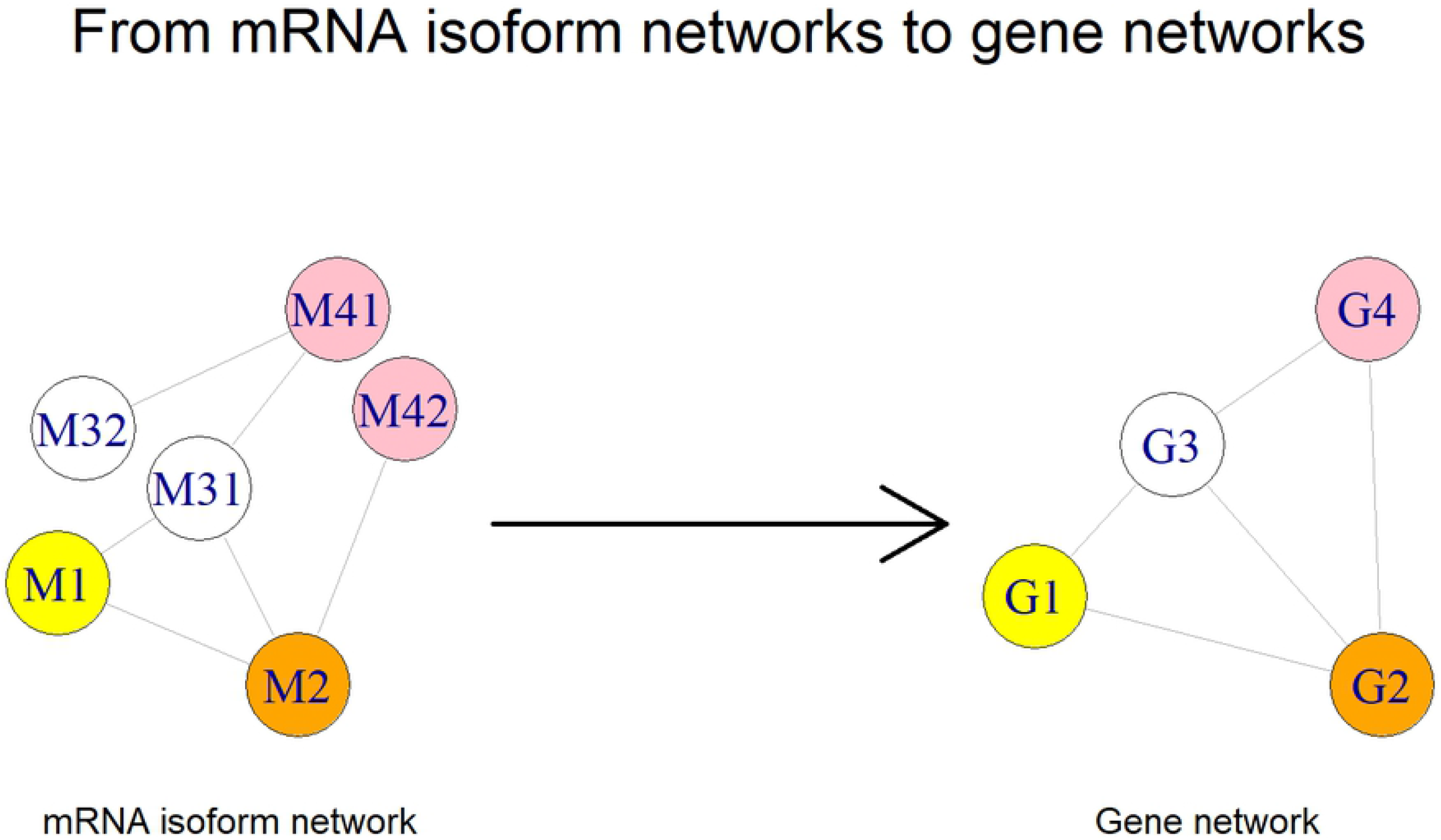
Constructing gene level networks from mRNA isoform networks. Shown here is the process by which we construct gene level networks using the tissue-specific functional mRNA isoform pair networks. All edges from the mRNA isoforms of the same gene in the mRNA isoform network are transferred to the single gene node in the gene level network. The gene and its mRNA isoforms have the same color.

### Tissue-specific network analysis

We use igraph [44] in R [32] for analyzing the graph properties of tissue-specific networks. We calculate basic statistics like number of nodes, number of edges, density, number of components, and size of the largest connected component for both mRNA isoform and gene level networks. Using the largest connected component for every network, we find central nodes (top 10%) using betweenness centrality and degree centrality. We also check the overlap between the central nodes as found using both centrality measures. The overlapping central gene nodes are further subjected to functional enrichment analysis.

In addition to calculating the global network properties, we also extract the mRNA isoforms, genes and gene pairs that are specific to a tissue and those that are shared by multiple tissues.

#### Functional enrichment analysis

We use the tissue-specific list of overlapping central gene nodes to perform functional enrichment analysis using the ReactomePA (version 1.26.0) and clusterProfiler (version 3.10.0) packages in R [32,45,46]. Enrichment is performed for Reactome pathways (version 66), KEGG pathways (release 88.2), GO biological process, GO molecular function and GO cellular components (GO data with a time stamp from the source of 10 October 2018 used by tools). In reactome data model, the core unit is a reaction while KEGG provides information about higher-level systemic functions of the cell and the organism. Due to the differences in the underlying data model and how pathways are defined, we perform enrichment analysis for both Reactome and KEGG pathways. We use a p-value cutoff of 0.05, false discovery rate control using Benjamini-Hochberg [47] with a cutoff of 0.05, minimum term size of 10, and maximum term size of 1000 for the enrichment analyses. We also remove redundant GO terms with a semantic similarity greater than 0.7 using the “Wang” measure [48] and keep the terms with most significant adjusted p-value. We further filter the GO terms to four levels [45,46] and plot only the top 5 most significant terms for every tissue. Neural tube was removed from the functional enrichment analysis because there was only 1 central gene.

### Model evaluation

#### Randomization experiments

To test the effect of randomization during the generation of training and testing datasets, we performed 1000 iterations of random training and testing dataset generation. In each iteration, we shuffle the combined functional and non-functional pairs, select 640,000 functional pairs and non-functional pairs respectively for the training dataset, select 160,000 functional and non-functional pairs respectively for the testing dataset, train a random forest model on the training dataset, use the trained model to make predictions on the testing dataset, and compute performance metrics. These datasets are referred to as “randomized datasets”.

To examine whether the random forest model learns genomic and sequence features that are predictive of functional mRNA isoform pairs, we perform a control experiment in which the functional and non-functional class labels are randomly shuffled to destroy the feature-class relationship in the original dataset. We perform 500 iterations of random training and testing dataset generation in which the functional and non-functional mRNA isoform pair class labels are shuffled. We train a random forest model on the class label shuffled training dataset, use the trained model to make predictions on the class label shuffled testing dataset and compute performance metrics. These datasets are referred to as the “class-label shuffled datasets”.

We also evaluate the impact of number of trees on the performance of the random forest model. For this, we use the following number of trees: 10, 20, 50, 100, 200, 500, 1000, 2000, and 5000. Again, we train one model with each of these number of trees using the training dataset and then evaluate the performance using the predictions on the testing dataset.

#### Using stratified cross-validation

We evaluate the performance of TENSION using a Stratified 10-Fold cross-validator. In terms of bias and variance, stratification, a sampling technique without replication and where class frequencies are preserved, is generally a better scheme as compared to regular cross-validation [49]. We use the original training data to create the 10-fold splits using StratifiedKFold function from Scikit-learn [42] which preserves the relative class frequency in each training and held out test fold. We then evaluate the performance of each fold by computing the AUROC and AUPRC using the predictions made on the held out test fold.

#### Validating predictions using new annotations

Because there is no gold standard dataset available for mRNA isoform level functions, we validate our predictions using the latest annotations from GO, KEGG pathways, BioCyc pathways, IntAct [26], BioGRID [38], APID [39], IID [40] and Mentha [41]. The new annotations (downloaded on 5 June 2018) were also processed as described in the “mRNA isoform level functional labels” section. Using our strategy to utilize the single isoform gene annotations for creating functional pairs, we found 284,916 functional pairs in the new annotations not present in our original functional pairs. Similarly, we found 112,827 non-functional pairs in the new annotations not present in our original non-functional pairs. We refer this new set of functional and non-functional mRNA isoform pairs as the “validation set”.

#### Validation of literature datasets

We also validate the predictions made by TENSION using two datasets from the literature: 1) a list of 20 ubiquitously expressed genes [50] and, 2) a list of 5035 genes that are expressed higher (expression fold change greater than 4 relative to all other tissues) in a specific tissue [51]. Only the tissues present in both TENSION and the transcriptomic BodyMap of mouse are selected for validation. We merge the three brain regions used in TENSION, forebrain, midbrain and hindbrain into a single brain entity for the analysis. Additionally, we removed the transcriptomic BodyMap of mouse genes that were not included in our initial 21,813 genes. This resulted in a final gene set of 1654 genes for the transcriptomic BodyMap of mouse. It is important to note that the above gene lists are based solely on the gene expression and do not necessarily translate to functionally enriched genes and as such we expect to find interactions involving these genes in multiple tissues.

#### Comparison with existing methods

To demonstrate the utility of using a simple supervised learning framework and improvements over previous methods for mRNA isoform functional network prediction, we compare TENSION with the Bayesian network based multi-instance learning model in [18]. We use our original training dataset with all 27 features to train the Bayesian network classifier and TENSION and make predictions on our original testing dataset. The output scores for mRNA isoform pairs in the original testing dataset from Bayesian network classifier and TENSION were used to compare the performance of the methods. We evaluate the performance of both the methods by computing the AUROC and AUPRC.

## Results

### A random forest model for functional network prediction

We use the mouse genome build GRCm38.p4 from NCBI in this study. After filtering the mRNA isoforms containing non-standard characters, less than 30 amino acid protein products and those missing either sequence or expression profile, we retained 2,874,753,225 mRNA isoform pairs. We have calculated 27 heterogeneous genomic and sequence-based features for all the mRNA isoform pairs (Table 1). Of these, we labelled 2,083,679 mRNA isoform pairs as functional pairs (positive) using the single mRNA isoform genes (described in methods section). And 818,071 mRNA isoform pairs as non-functional pairs (negative) by using the “NOT” annotation tag in the GO annotations (described in methods section). These functional and non-functional mRNA isoform pairs are used to train and develop random forest models for predicting mouse mRNA isoform level functional networks. The predictions made by random forest have an associated probability score which measures the strength of mRNA isoform interactions.

#### Randomization experiments

Randomization experiments test the effect of selecting functional and non-functional pairs when generating training and testing datasets. Fig 4 shows that there is very little to no variance in the performance of randomized datasets. Therefore, we generate one final training and testing dataset (“original datasets”) by randomly selecting functional and nonfunctional pairs and use it to generate the final functional network prediction models.

**Fig 4.**
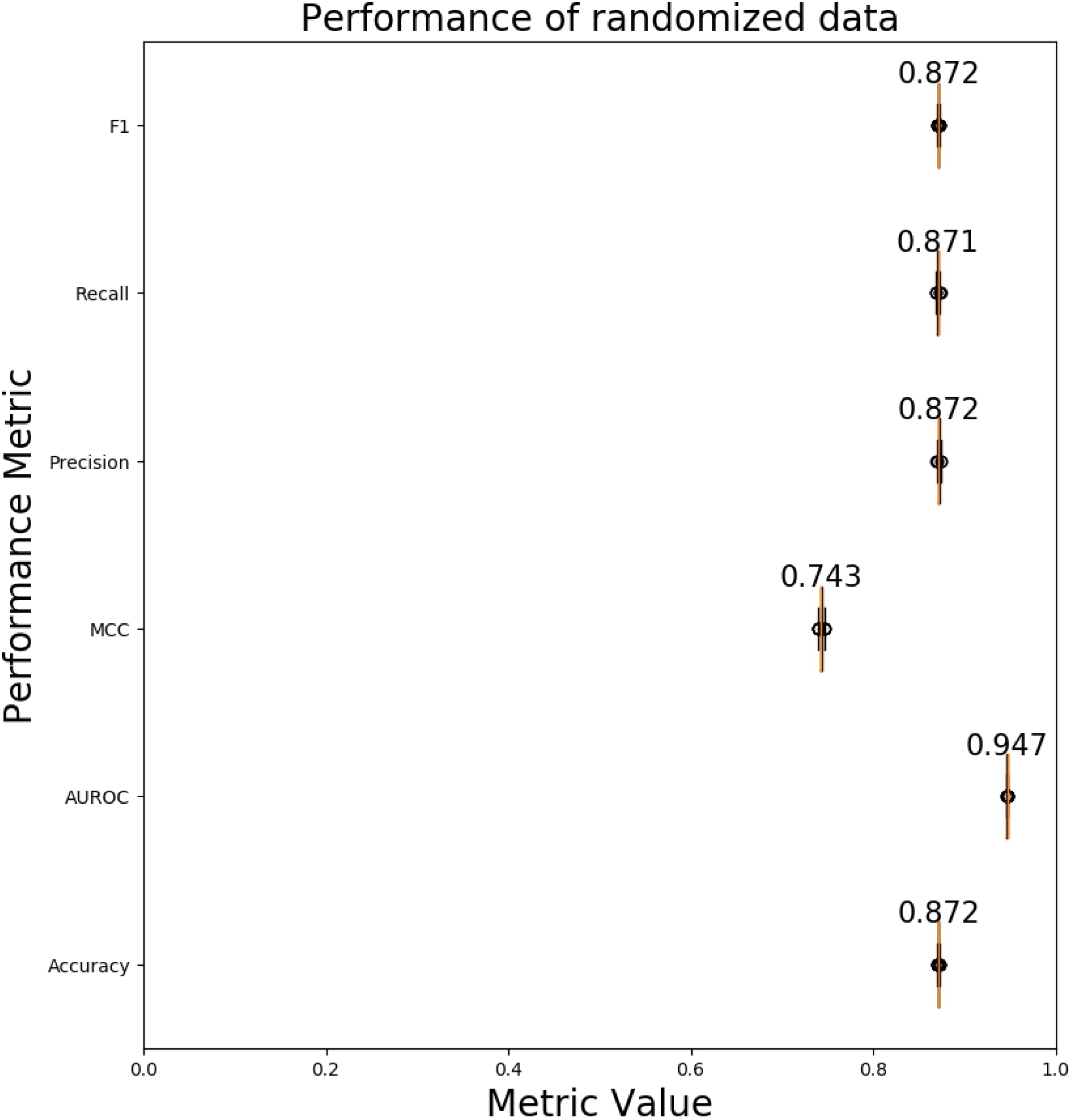
Performance evaluation on randomized datasets. A boxplot of various performance evaluation metrics calculated using 1000 randomized datasets. The median value is shown for the performance metrics. The width of the boxes along the x-axis represent the variability in the value of the performance metric across 1000 randomized datasets. Higher metric value and smaller box width is better. Abbreviations - AUROC: Area Under the Receiver Operating Characteristic Curve; MCC: Matthews Correlation Coefficient

To help us identify if TENSION is actually learning from the data and not just making random predictions, we estimate the performance of the random forest model on the class-label shuffled datasets. The AUROC obtained on the class-label shuffled datasets is 0.5 (as compared with 0.947 on the original testing dataset) indicating that our functional network prediction model performs significantly better than random predictions (Fig 5).

**Fig 5.**
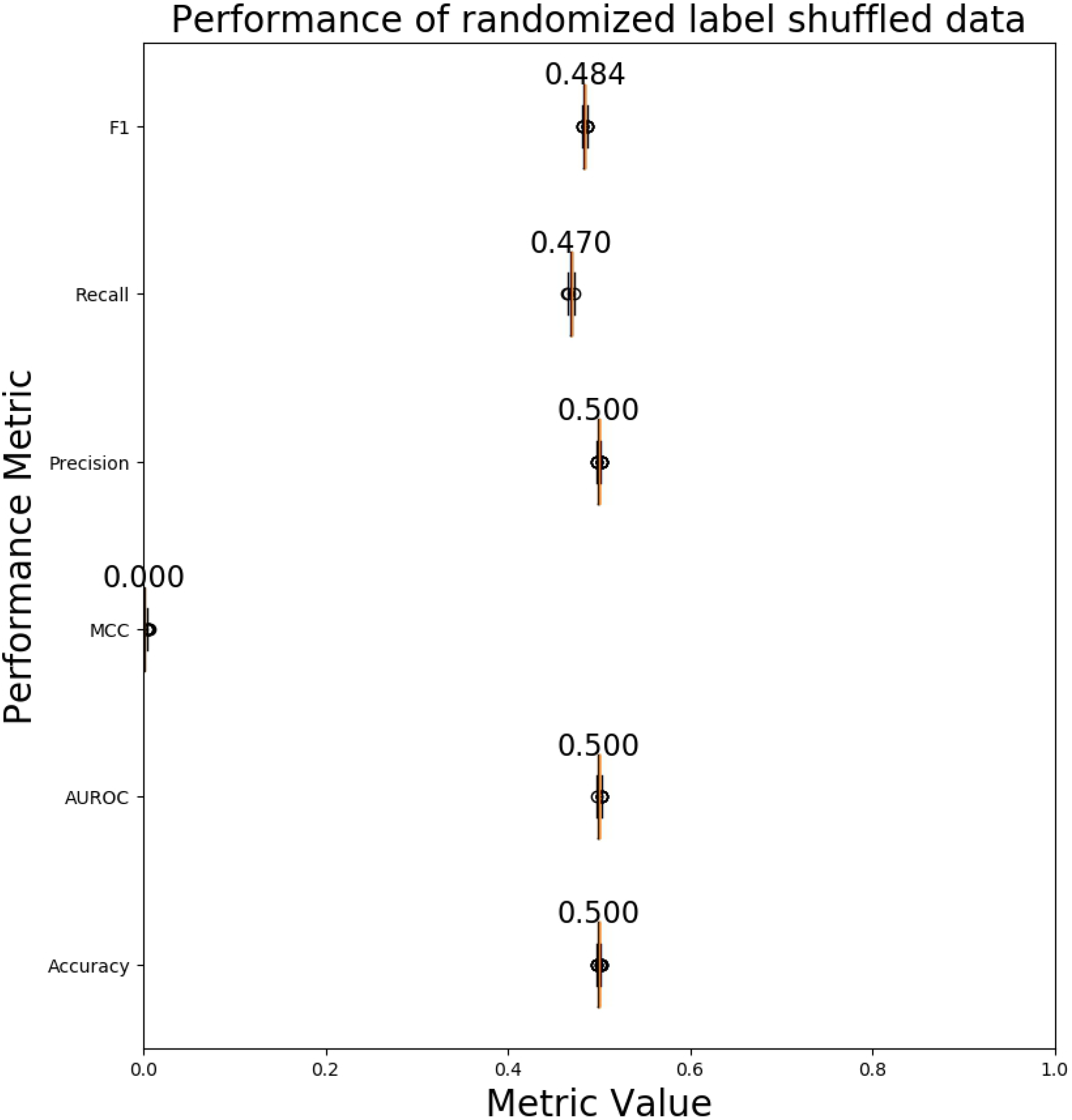
Performance evaluation on label shuffled datasets. A boxplot of performance evaluation metrics calculated using 500 label shuffled datasets. The functional and non-functional labels for mRNA isoform pairs are randomly shuffled while still maintaining the class distribution (equal functional/non-functional pairs). The median value is shown for the performance metrics. The width of the boxes along the x-axis represent the variability in the value of the performance metric across 500 label shuffled datasets. Higher metric value and smaller box width is better. The performance of a model which makes random guesses is about 0.5 (or 0 for MCC because it ranges from −1 to 1). Abbreviations - AUROC: Area Under the Receiver Operating Characteristic Curve; MCC: Matthews Correlation Coefficient.

#### Performance evaluation

We evaluate the performance of TENSION when using all 27 features from the predictions on the original testing dataset. We first evaluate the impact of number of trees on the performance of random forest model. It can be seen in Fig S1 that there is very little improvement in the performance of the model after 100 trees. To reduce computational complexity without sacrificing the performance while making predictions for all 2.9 billion mRNA isoform pairs, we use 100 trees in our final models. On the original testing dataset, we obtain a high correlation as seen in Table 2 and Fig S2 suggesting a highly accurate model.

**Table 2.**
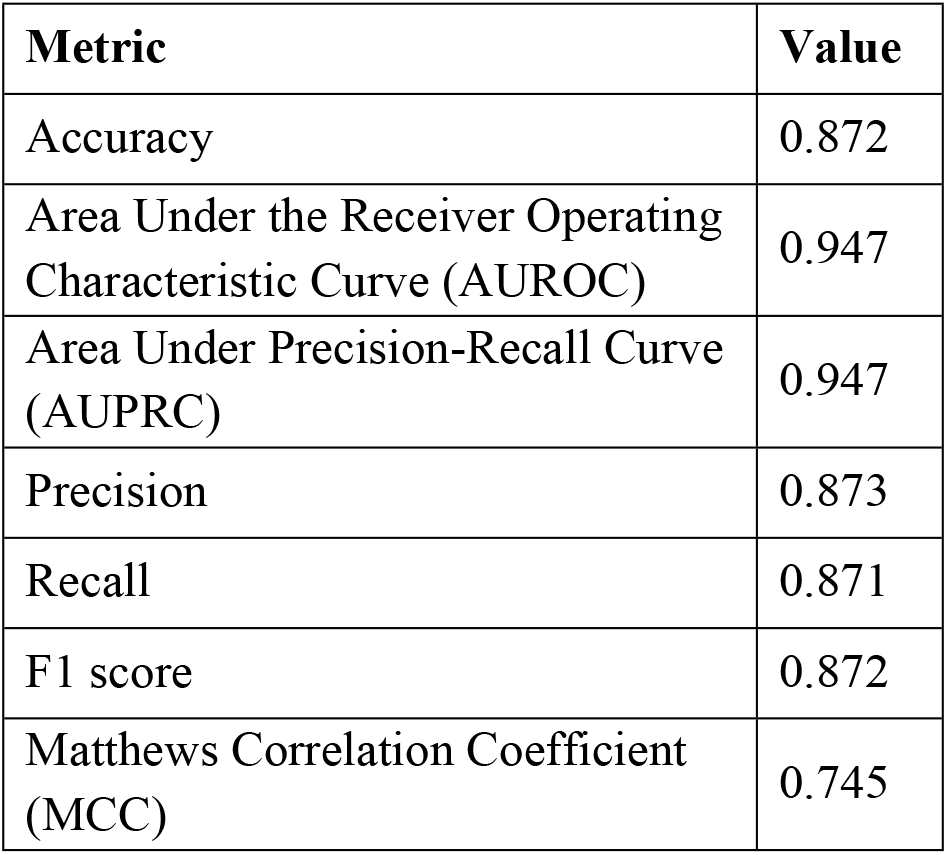
Prediction performance metrics for TENSION on the original testing dataset with all 27 features.

| Metric | Value |
| --- | --- |
| Accuracy | 0.872 |
| Area Under the Receiver Operating Characteristic Curve (AUROC) | 0.947 |
| Area Under Precision-Recall Curve (AUPRC) | 0.947 |
| Precision | 0.873 |
| Recall | 0.871 |
| F1 score | 0.872 |
| Matthews Correlation Coefficient (MCC) | 0.745 |

#### Evaluation by stratified cross-validation

In addition to evaluating the performance of our random forest on a held-out test set, we also perform stratified 10-fold cross validation. The AUROC and AUPRC curves for each fold are shown in Fig 6. We see that there is very little variance in the results of each fold. The results are also very close to those obtained on the original testing dataset (S2 Fig). The results of stratified cross-validation emphasize the robustness of TENSION.

**Fig 6.**
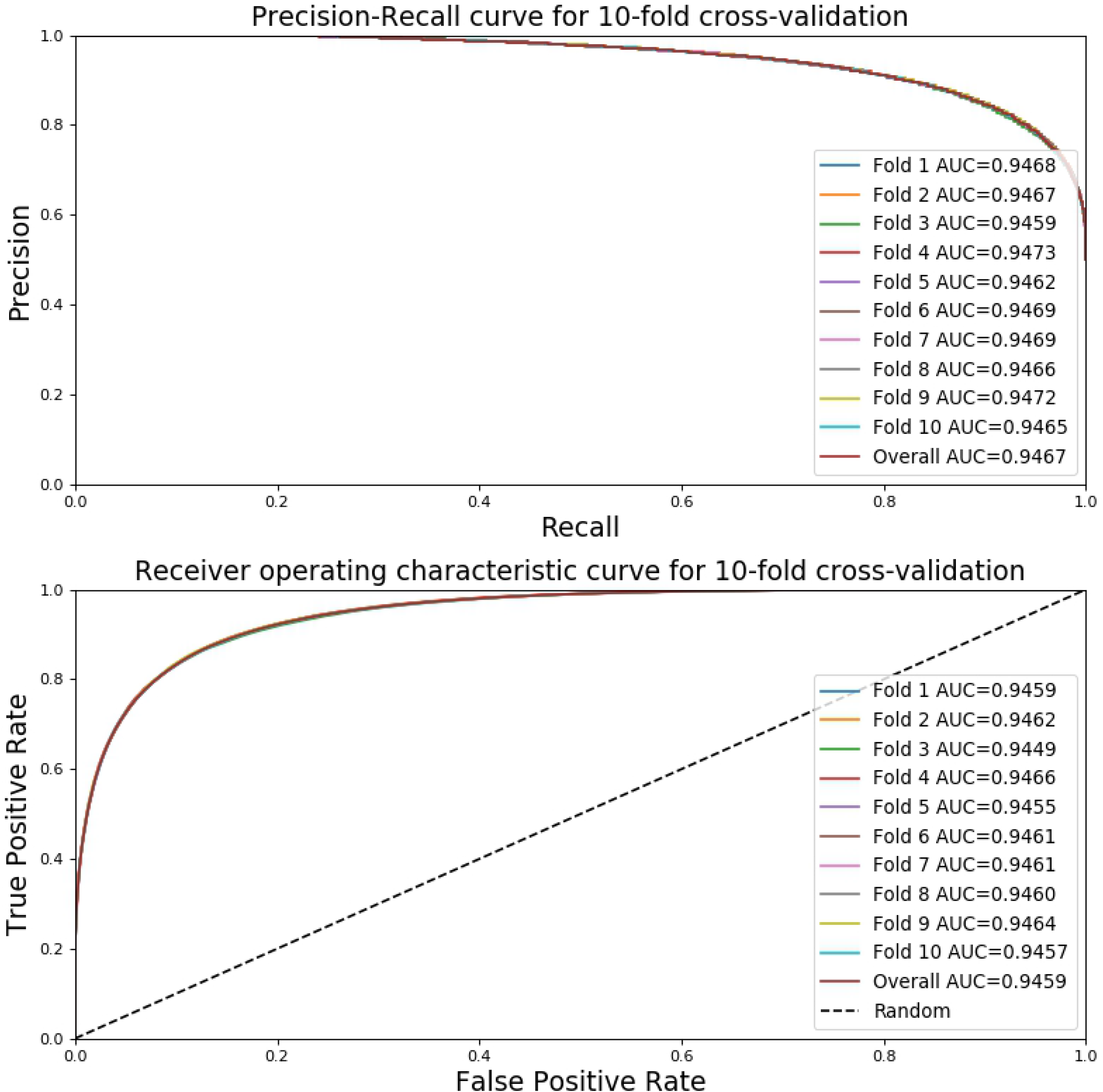
Performance evaluation by 10-fold stratified cross-validation. The precision-recall and receiver operating characteristic curve for all 10 folds of the stratified cross-validation. Note that the performance is virtually identical for all folds suggesting the robustness of TENSION. A model with area under the curve closer to 1 is better while a model with an area under the curve of 0.5 is equivalent to making random guess. Abbreviations - AUC: Area Under the Curve.

#### Validating predictions using new annotations

After processing the new GO annotations, pathway, and PPIs data, we learned a new set of 397,743 previously unknown mRNA isoform pairs. Of these, we labelled 284,916 as functional and 112,827 as non-functional mRNA isoform pairs. Using all 27 features, TENSION correctly classified 315,844 (out of 397,743) mRNA isoforms pairs at an overall accuracy of 79.4%. The true positives, true negatives, false positives, and false negatives collectively represented by a confusion matrix are presented in Table 3. Since the distribution of functional and non-functional mRNA isoform pairs in the validation set is imbalanced, we also access the performance of our classifier by computing the AUPRC and AUROC. We observe an AUPRC of 0.926 and an AUROC of 0.855 (Fig 7). In addition to these curves, we also calculate the Precision (0.885), Recall (0.819), F1 score (0.851) and MCC (0.524). These are much higher than random predictions shown in Fig 5 suggesting that TENSION performs better than random guessing and is also able to predict potential functional and nonfunctional mRNA isoform pairs accurately.

**Table 3.** Confusion matrix for predictions on validation set.

| **True Label ↓** | **Predicted Label** |  |  |
| --- | --- | --- | --- |
|  | Functional | Non-Functional |  |
| Functional | 233434 | 51482 | **284916 (81.9%)** |
| Non-Functional | 30417 | 82410 | **112827 (73%)** |
|  | **263851** | **133892** |  |

**Fig 7.**
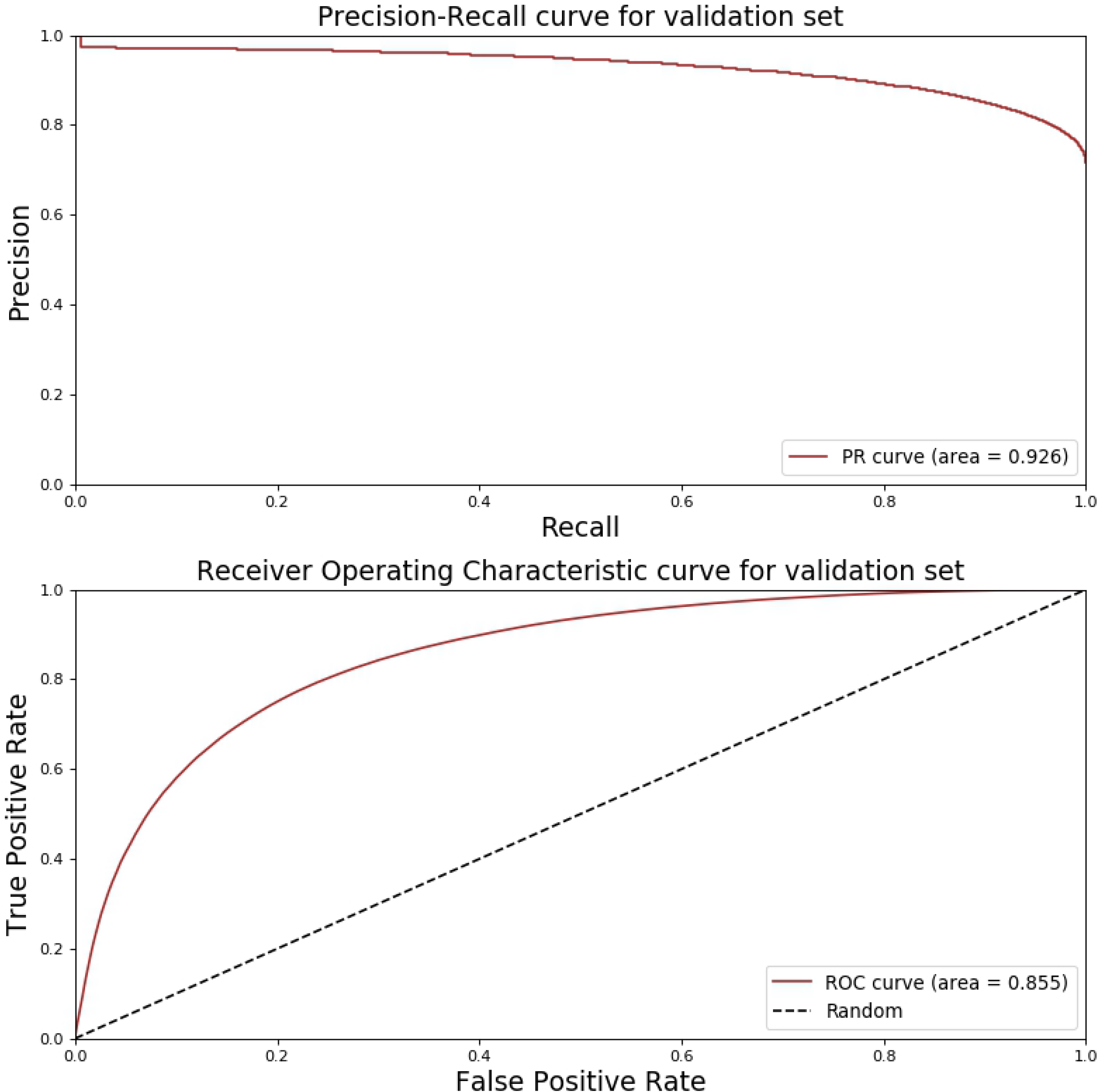
Performance evaluation on validation dataset. The precision-recall and receiver operating characteristic curve for predictions on the validation dataset. The validation dataset is constructed by using the later version of gene ontology annotations, pathways and protein-protein interactions than those used for our original mRNA isoform level label generation. A model with area under the curve closer to 1 is better while a model with an area under the curve of 0.5 is equivalent to making random guess. Abbreviations - PR: Precision-Recall; ROC: Receiver Operating Characteristic.

Of these new mRNA isoform pairs, 8200 are predicted as tissue-specific functional mRNA isoform pairs. However, the annotations in GO, KEGG, BioCyc, and PPI databases do not store tissue information, so we cannot validate the tissue specificity of these predictions.

#### Comparison with existing methods

We compare the performance of TENSION when using all 27 features with that of the Bayesian network based MIL method [18]. The default parameters are used for the Bayesian network-based MIL method. We use our original training dataset to train the Bayesian network-based MIL method and TENSION and then make predictions on our original testing dataset. We calculate the AUROC and AUPRC using these predictions for both models to compare their performance. The functional mRNA isoform pairs are derived from single mRNA producing genes co-annotated to GO biological process, pathways or PPIs whereas the nonfunctional mRNA isoform pairs are constructed by using the ‘NOT’ tagged GO biological process annotations.

The Bayesian network based MIL method achieves an AUROC of 0.761 (Fig 8) which is higher than the original AUROC value of 0.656 reported in the original study [18]. TENSION achieves significantly higher AUROC of 0.947. Similarly, TENSION achieves significantly higher AUPRC of 0.947 as compared to Bayesian network based MIL method’s AUPRC of 0.757 (Fig 8). The significantly higher AUROC and AUPRC values of TENSION highlights the importance of using a simple supervised learning framework and improvements over the more complex MIL-based methods for mRNA isoform functional network prediction. It should be noted that the MIL-based method was originally developed using different set of features, however, for the purpose of comparison we have used the same training and testing datasets for both methods. The improved performance of Bayesian network based MIL method on our dataset also highlights the significance of mRNA isoform level functional label and feature generation in TENSION.

**Fig 8.**
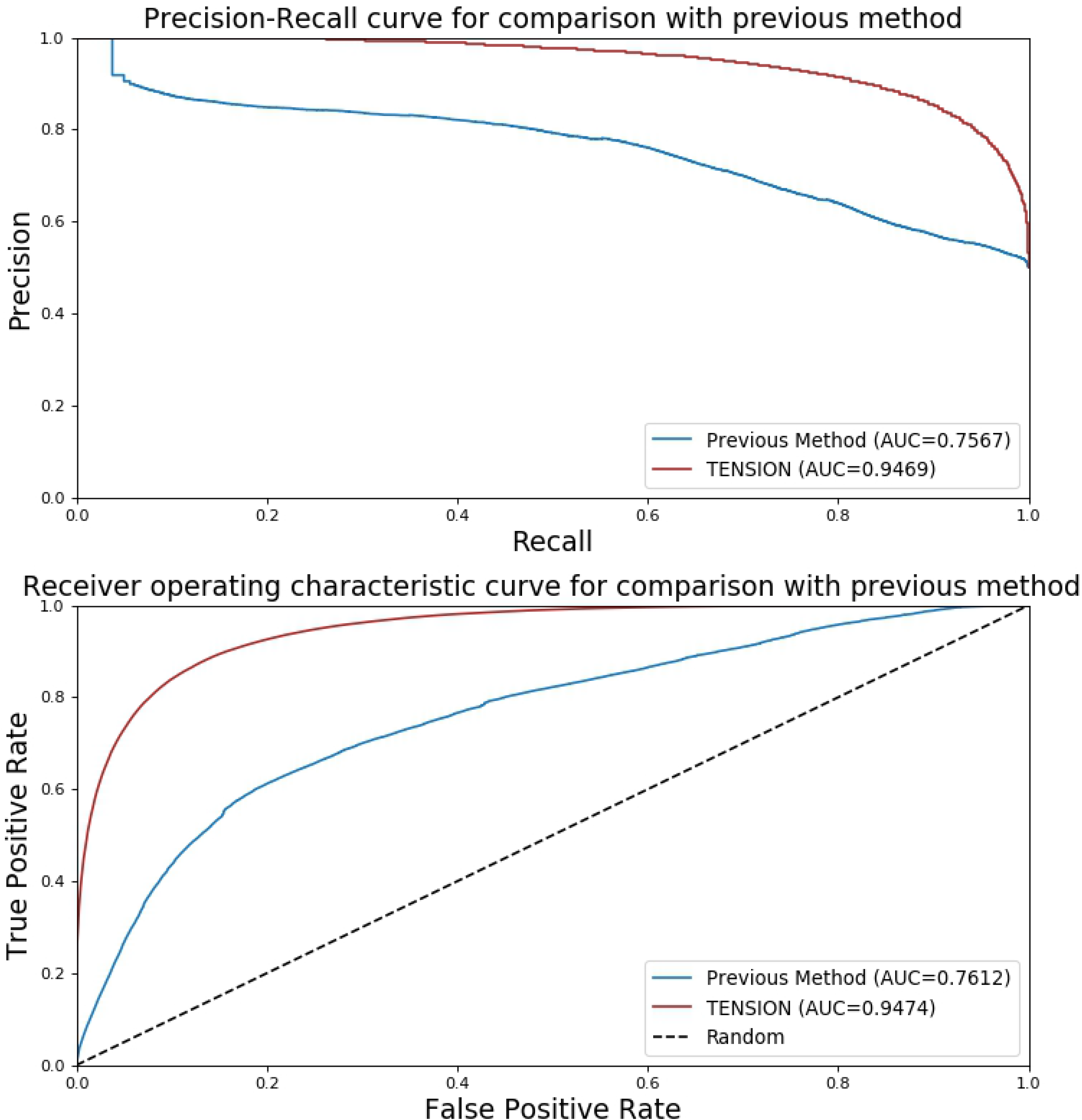
Performance comparison with Bayesian network based multi-instance learning method. The precision-recall and receiver operating characteristic curve for performance comparison of TENSION with previously published Bayesian network based multi-instance learning method. The original training dataset was used to train both models and performance was calculated using the predictions made on the original testing dataset. Higher values of area under the curve correspond to more accurate predictions. Abbreviations - AUC: Area Under the Curve.

### Tissue-specific networks

#### Tissue-specific functional mRNA isoform pair networks

As shown in Fig 2, to build the tissue-specific mRNA isoform level functional networks, we assume that, for a tissue ***i***, if an mRNA isoform pair is predicted to be functional (positive) using all 27 features, but the prediction after removing the tissue ***i***-specific feature is non-functional (negative), the mRNA isoform pair is only functional under tissue ***i***. The strength of mRNA isoform interactions is measured by the probability score predicted by random forest. To remove noise, low confidence predictions and organism-wide reference mRNA isoform pairs from tissue-specific functional networks, we only consider the mRNA isoform pairs which have a random forest predicted probability score ≥ **0.6** when using all 27 features and a probability score ≤ **0.4** after removing the tissue derived RNA-Seq feature. For the tissue specific functional networks, a lower probability score corresponds to higher strength of mRNA isoform pair to be involved in the same GO biological process or pathway. A summary of all 17 tissue-specific mRNA isoform functional networks as obtained after applying the above filtering criteria is provided in Table 4.

**Table 4.**
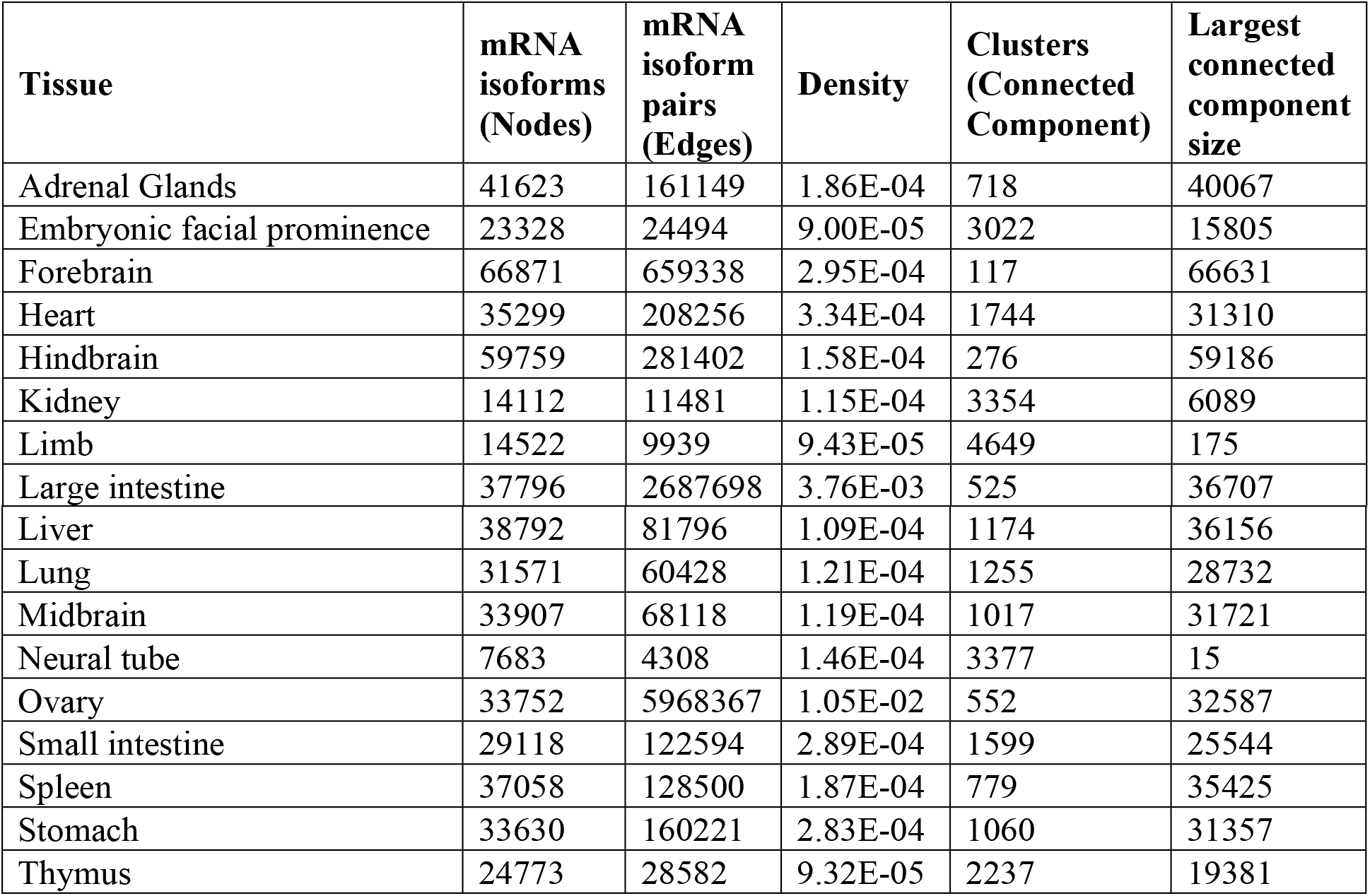
Summary statistics for mRNA isoform level single tissue functional networks.

| Tissue | mRNA isoforms (Nodes) | mRNA isoform pairs (Edges) | Density | Clusters (Connected Component) | Largest connected component size |
| --- | --- | --- | --- | --- | --- |
| Adrenal Glands | 41623 | 161149 | 1.86E-04 | 718 | 40067 |
| Embryonic facial prominence | 23328 | 24494 | 9.00E-05 | 3022 | 15805 |
| Forebrain | 66871 | 659338 | 2.95E-04 | 117 | 66631 |
| Heart | 35299 | 208256 | 3.34E-04 | 1744 | 31310 |
| Hindbrain | 59759 | 281402 | 1.58E-04 | 276 | 59186 |
| Kidney | 14112 | 11481 | 1.15E-04 | 3354 | 6089 |
| Limb | 14522 | 9939 | 9.43E-05 | 4649 | 175 |
| Large intestine | 37796 | 2687698 | 3.76E-03 | 525 | 36707 |
| Liver | 38792 | 81796 | 1.09E-04 | 1174 | 36156 |
| Lung | 31571 | 60428 | 1.21E-04 | 1255 | 28732 |
| Midbrain | 33907 | 68118 | 1.19E-04 | 1017 | 31721 |
| Neural tube | 7683 | 4308 | 1.46E-04 | 3377 | 15 |
| Ovary | 33752 | 5968367 | 1.05E-02 | 552 | 32587 |
| Small intestine | 29118 | 122594 | 2.89E-04 | 1599 | 25544 |
| Spleen | 37058 | 128500 | 1.87E-04 | 779 | 35425 |
| Stomach | 33630 | 160221 | 2.83E-04 | 1060 | 31357 |
| Thymus | 24773 | 28582 | 9.32E-05 | 2237 | 19381 |

The tissue-specific functional networks identify around 10.6 million tissue-specific functional mRNA isoform pairs (0.37% of all possible mRNA isoform pairs). The density of tissue-specific functional networks is in the order of 10^−2^ – 10^−5^ and most networks are very sparse. The number of tissue-specific functional mRNA isoform pairs vary greatly across the tissues, from few thousands in limb and neural tube to few million in large intestine and ovary (Table 4). All these mRNA isoform pairs are present in only one tissue. Table 4 shows the number of functional mRNA isoform pairs identified as single tissue-specific in each of the 17 tissues.

All tissues have many connected components (Table 4). Limb, neural tube and kidney have less than 50% mRNA isoform nodes in their largest connected component, whereas some others like hindbrain, large intestine, ovary, and forebrain etc. have over 90%. These differences in the size of networks, mRNA isoforms involved and the network structures highlight the differences in tissue-level biological processes as evident by the differences in the enriched pathways and gene ontology terms (discussed later).

To highlight the differences that arise when analyzing functional networks at the mRNA isoform and gene level and because all functional enrichment tools are built for analyzing genes, we also compress the mRNA isoform level networks to gene level networks. In the gene level networks, all mRNA isoform nodes of the same gene are combined into a single gene node. Table 5 provides a summary of the gene level networks for all 17 tissues. We identified around 7.79 million unique gene pairs (3.27% of all possible gene pairs) using these tissue level gene functional networks. It was recently observed in mouse and humans that testis and ovary express the highest number of genes whereas brain and liver express the highest number of tissue enriched genes under normal conditions [51,52]. This is also reflected in our gene level networks (Table 5) where ovary, hindbrain and forebrain networks have the largest number of edges (gene pairs) and nodes (genes).

While the majority of gene pairs are present in only one tissue level gene functional networks (98% of identified gene pairs; Table 5), a small fraction (2% of identified gene pairs; Table 5 and Fig 9) is present in at least two tissue level gene functional networks. Although the gene pairs are shared between tissues, the mRNA isoform pairs resulting from these gene pairs are specific to only one tissue. This highlights that different mRNA isoforms of the same gene can have different functional partners across tissues.

**Table 5.** Summary statistics for gene level functional networks.

| Tissue | Genes (Edges) | Gene pairs (Edges) | Tissue specific gene pairs | Density | Clusters | Largest connected component size | Gene pairs shared with other tissues |
| --- | --- | --- | --- | --- | --- | --- | --- |
| Adrenal gland | 17171 | 146413 | 137526 | 9.33E-04 | 153 | 16953 | 8887 |
| Embryonic facial prominence | 13058 | 24183 | 22346 | 2.62E-04 | 610 | 12001 | 1837 |
| Forebrain | 19979 | 509765 | 485982 | 2.44E-03 | 22 | 19944 | 23783 |
| Heart | 15874 | 189042 | 178026 | 1.41E-03 | 274 | 15452 | 11016 |
| Hindbrain | 19546 | 239309 | 228381 | 1.20E-03 | 44 | 19491 | 10928 |
| Kidney | 9672 | 11226 | 10413 | 2.23E-04 | 1236 | 7059 | 813 |
| Limb | 9619 | 9677 | 9139 | 1.98E-04 | 1300 | 6689 | 538 |
| Large intestine | 14981 | 1999465 | 1893127 | 1.69E-02 | 173 | 14648 | 106338 |
| Liver | 17420 | 77521 | 72872 | 4.80E-04 | 147 | 17216 | 4649 |
| Lung | 14784 | 56739 | 52527 | 4.81E-04 | 255 | 14402 | 4212 |
| Midbrain | 15581 | 62722 | 59576 | 4.91E-04 | 211 | 15265 | 3146 |
| Neural tube | 6057 | 4278 | 3978 | 2.17E-04 | 2083 | 176 | 300 |
| Ovary | 14613 | 4207053 | 4079552 | 3.82E-02 | 196 | 14221 | 127501 |
| Small intestine | 14254 | 116777 | 110355 | 1.09E-03 | 345 | 13676 | 6422 |
| Spleen | 15320 | 119865 | 110709 | 9.43E-04 | 234 | 14962 | 9156 |
| Stomach | 14991 | 152581 | 142860 | 1.27E-03 | 238 | 14589 | 9721 |
| Thymus | 13145 | 27451 | 25378 | 2.94E-04 | 458 | 12374 | 2073 |

**Fig 9.**
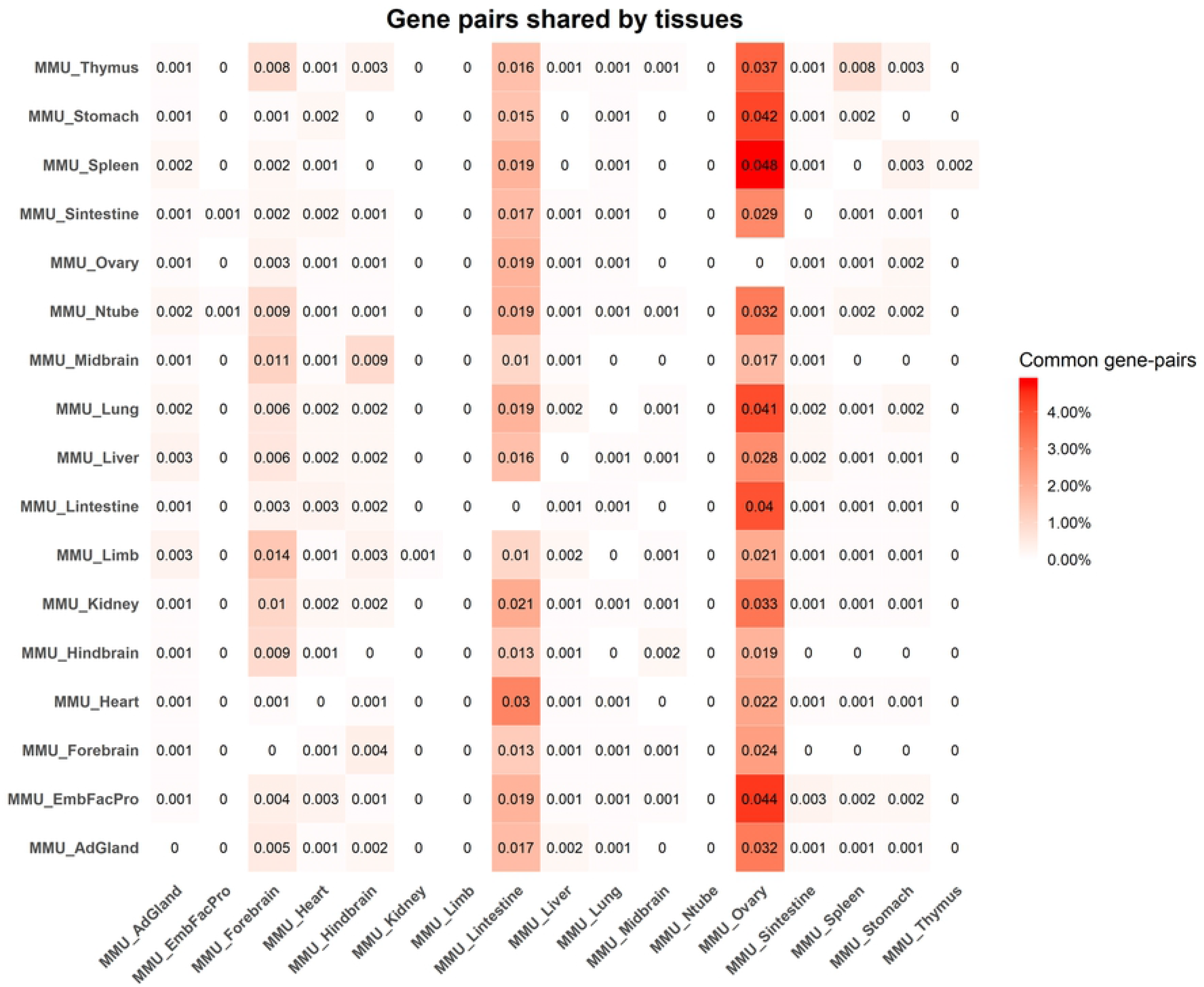
Fraction of gene pairs shared between tissues. The heatmap represents the fraction of gene pairs shared between two tissues. The numbers shown in the heatmap are not symmetric because the fraction is weighted by total gene pairs in that row’s tissue. The fraction is weighted by the total number of pairs in the tissue specified on row. For instance, spleen shares 4.8% of all gene pairs present in the spleen network with ovary. Darker shades refer to higher fractions of shared gene pairs. The numbers in the heatmap should be interpreted as reading a matrix rowwise. Abbreviations - AdGland: Adrenal glands; EmbFacPro: Embryonic facial prominence; Sintestine: Small intestine; Lintestine: Large intestine.

Shared gene-pairs may indicate shared processes between tissues. The spleen and embryonic facial prominence share the highest fraction of gene pairs (about 7.6% of gene pairs; Table 5) with other tissues, while ovary shares the lowest fraction (3% of all ovary gene pairs; Table 5). The composition of gene pairs shared between the tissue level functional networks is quite complex and is shown in Fig 9. Upon further investigation, we find that the spleen network shares 4.8% of its gene pairs with ovary network while ovary network shares only 0.1% of its gene pairs with the spleen network. We also find that thymus shares about 3.7% of its gene pairs with ovary, supporting the notion that thymus is necessary for normal ovarian development and function after the neonatal period [53,54]. These findings further justify the importance of our networks in characterizing tissue level processes.

Like the mRNA isoform networks, the gene-level neural tube network contains only 2.9% of genes in its largest connected components (Tables 4 and 5). All other gene-level tissue networks have a very high fraction of genes and gene pairs in the largest connected components (Table 5).

#### Central genes in tissue-specific functional networks have tissue related characteristics

The central genes identified in our tissue-specific networks are enriched in tissue related GO terms and pathways (Figs 10 and 11). The central genes in the heart specific gene network are significantly enriched in transmembrane transporter activity, vitamin binding, complement and coagulation cascades etc. (Figs 10 and 11). Supplementation of several vitamins such as Vitamin B6, Vitamin D, Vitamin E, and folate etc. are linked to reduced risk of cardiovascular diseases [55–59]. The serine proteinase cascades of the coagulation and the complement systems have been associated with functions of the cardiovascular and immune systems [60].

**Fig 10.**
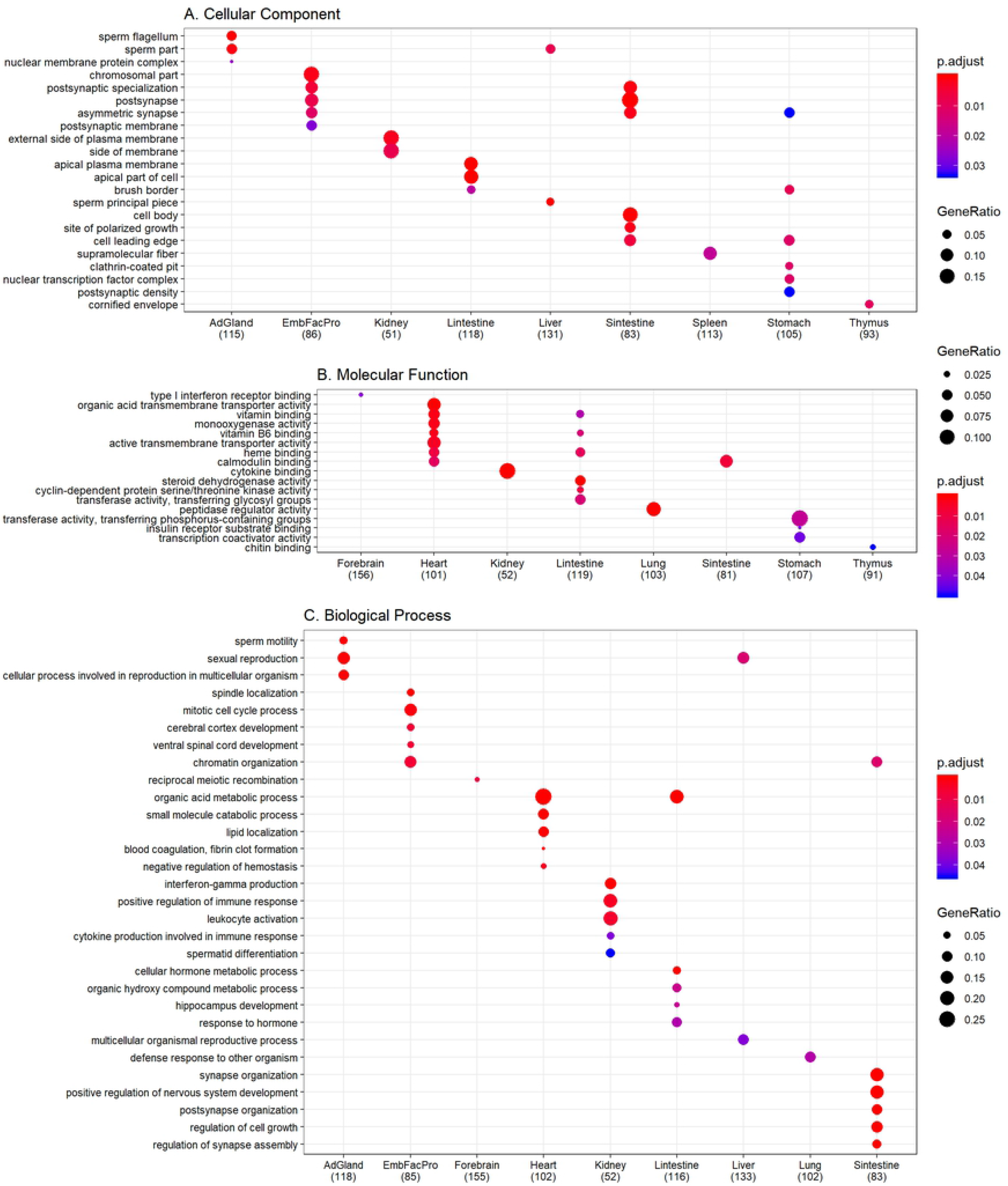
Gene ontology functional enrichment. Since the functional annotations are at the gene level, we use the central genes identified by both betweenness centrality (top 10%) and degree centrality (top 10%) to perform gene ontology enrichment. Only the top 5 terms for every tissue are shown here. The dot size represents the ratio of genes present in our central genes annotated to a gene ontology term to genes present in our central network. The color signifies the value of adjusted p-value from false discovery rate control using Benjamini-Hochberg, with lower adjusted p-values shown in darker intensities of red. **A.** Enrichment for cellular component aspect of gene ontology. **B.** Enrichment for molecular function aspect of gene ontology. **C.** Enrichment for biological process aspect of gene ontology. Abbreviations – AdGland: Adrenal glands; EmbFacPro: Embryonic facial prominence; Sintestine: Small intestine; Lintestine: Large intestine.

**Fig 11.**
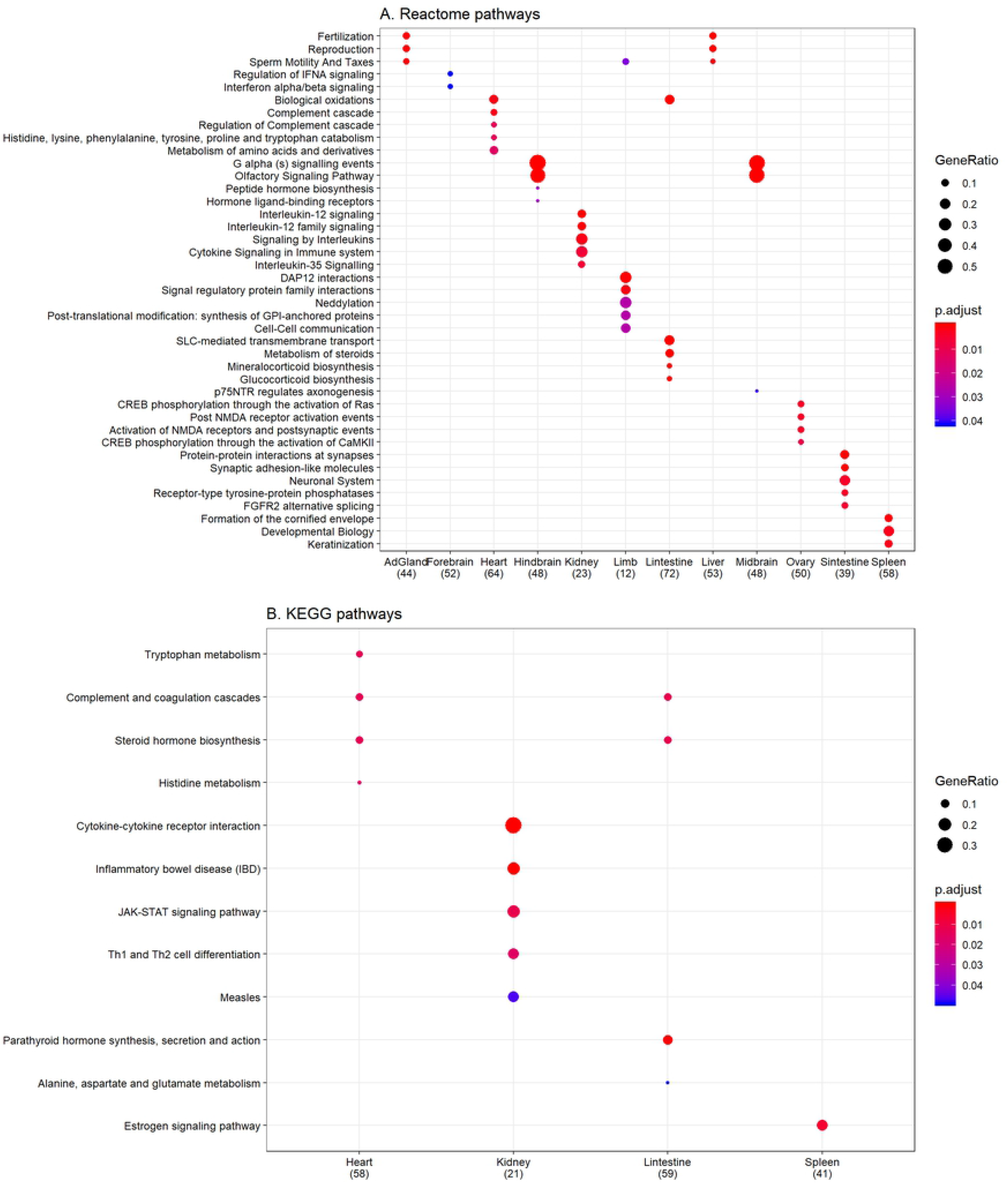
Pathway enrichment analysis. We use the central genes identified by both betweenness centrality (top 10%) and degree centrality (top 10%) to perform pathway enrichment. Only the top 5 pathways for every tissue are shown here. The dot size represents the ratio of genes present in our central genes annotated to a pathway to genes present in our central network. The color signifies the value of adjusted p-value from false discovery rate control using Benjamini-Hochberg, with lower adjusted p-values shown in darker intensities of red. **A.** Enrichment for reactome pathways. **B.** Enrichment for KEGG pathways. Abbreviations - KEGG: Kyoto Encyclopedia of Genes and Genomes; AdGland: Adrenal glands; Sintestine: Small intestine; Lintestine: Large intestine.

Several important renal processes such as JAK-STAT signaling pathway, cytokine signaling in immune system, cytokine-cytokine receptor interaction, and signaling by interleukins etc. are enriched in the central genes of the kidney specific gene network (Figs 10 and 11). Defects in these processes and pathways have been linked to several renal disorders and related co-morbidities [61–64]. Genes in the kidney network are also enriched for interferon-gamma production and inflammatory bowel disease (IBD; Figs 10 and 11). In IBD, interferon-gamma negatively regulates the Na^+^/Ca^2+^ exchanger 1 (NCX1) -mediated renal Ca^2+^ absorption contributing to IBD-associated loss of bone mineral density and altered Ca^2+^ homeostasis [65].

The large intestine has specific and efficient carrier mediated transporter mechanisms for the absorption of several water soluble vitamins (pantothenic acid, biotin, thiamin, riboflavin and folate) [66]. These vitamins are essential for several biological processes and their enrichment in large intestine specific gene network only seems natural (Figs 10 and 11). The *brain-in-the-gut* or the enteric nervous system (ENS) is the largest component of the autonomous nervous system [67–69]. The small intestine ENS is equipped to perform functions relating to inflammation, digestion, secretion and motility among others [67–69]. The identification of several neuronal terms for central genes in the small intestine network is in line with such literature findings (Figs 10 and 11) [67–69].

Fertility and energy metabolism are reciprocally regulated and tightly linked in female animals and this relation has been conserved throughout evolution [70–72]. Metabolic disorders such as those of the liver can lead to changes in reproductive functions and vice-versa [70–72]. It was recently proposed that in case of protein scarcity, the estrous cycle is blocked and the liver acts as a critical mediator of reproductive and energetic functions [70–72]. The enrichment of several reproduction and fertility related terms in our liver specific network also point towards such observations (Figs 10 and 11).

We also find significantly enriched tissue related process terms for other tissues such as spleen, ovary, adrenal glands and limb etc. (Figs 10 and 11). However, the tissue specific central genes do not always lead to significantly enriched terms.

The identification of tissue related biological processes via the central genes highlights that TENSION can correctly capture the tissue-specific functional mRNA isoform pairs produced by genes involved in tissue related functions. We can identify the specific mRNA isoforms of these genes by looking back at the mRNA isoform level tissue networks. Finding the specific mRNA isoforms responsible for these processes should provide a significant clue towards understanding of developmental and molecular processes of diseases and biological functions.

#### Tissue-specific non-functional mRNA isoform pair networks

To build the tissue-specific mRNA isoform level non-functional networks, we assume that, for a tissue ***i***, if an mRNA isoform pair is predicted to be non-functional (negative) using all 27 features but the prediction after removing the tissue ***i***-specific feature changes to functional (positive), the mRNA isoform pair is only non-functional under tissue ***i*** (Fig 2). To remove noise and low confidence predictions in tissue-specific non-functional mRNA isoform networks, we only consider the mRNA isoform pairs which have a random forest predicted probability score of ≤ **0.4** when using all 27 features and a probability score of ≥ **0.6** after removing the tissue derived RNA-Seq feature. Higher probability score reflects stronger tissue-specific non-functional mRNA isoform pair. A summary of all 17 tissue-specific mRNA isoform level non-functional networks as obtained after applying the above filtering criteria is provided in Tables 6.

**Table 6.** Summary statistics for single tissue mRNA isoform level non-functional networks.

| **Tissue** | **mRNA isoforms (Nodes)** | **mRNA isoform pairs (Edges)** | **Density** | **Clusters** | **Largest connected component size** |
| --- | --- | --- | --- | --- | --- |
| Adrenal glands | 35368 | 213426 | 3.41E-04 | 3043 | 28354 |
| Embryonic facial prominence | 15194 | 8856 | 7.67E-05 | 6341 | 22 |
| Forebrain | 63151 | 1407212 | 7.06E-04 | 98 | 62947 |
| Heart | 30603 | 26859 | 5.74E-05 | 5817 | 15456 |
| Hindbrain | 62114 | 509639 | 2.64E-04 | 219 | 61653 |
| Kidney | 16503 | 10427 | 7.66E-05 | 6095 | 1247 |
| Large intestine | 47731 | 89559 | 7.86E-05 | 1357 | 44697 |
| Limb | 48172 | 112519 | 9.70E-05 | 754 | 46585 |
| Liver | 45339 | 114783 | 1.12E-04 | 1473 | 42159 |
| Lung | 14585 | 8380 | 7.88E-05 | 6213 | 28 |
| Midbrain | 23621 | 15607 | 5.59E-05 | 8036 | 163 |
| Neural tube | 18465 | 11510 | 6.75E-05 | 6958 | 40 |
| Ovary | 55783 | 799229 | 5.14E-04 | 526 | 54638 |
| Small intestine | 40655 | 99825 | 1.21E-04 | 1611 | 37148 |
| Spleen | 23949 | 37927 | 1.32E-04 | 5479 | 10275 |
| Stomach | 26541 | 23988 | 6.81E-05 | 4982 | 14012 |
| Thymus | 21565 | 22458 | 9.66E-05 | 6308 | 5189 |

Using these tissue-specific mRNA isoform level non-functional networks we identified around 3.5 million tissue-specific non-functional mRNA isoform pairs (0.12% of all possible mRNA isoform pairs). The tissue-specific non-functional networks are also sparse with density in the order of 10^−3^ – 10^−5^. The number of tissue-specific non-functional mRNA isoform pairs also vary greatly across the tissues. For instance, forebrain has a very high number of 1.4 million (40% of all tissue-specific non-functional mRNA isoform pairs) non-functional mRNA isoform pairs. All these mRNA isoform pairs are specifically non-functional in only one tissue.

Similar to the functional networks, we also compress the non-functional mRNA isoform networks to gene level non-functional networks. In the gene level networks, all mRNA isoform nodes of the same gene and their edges are combined into a single gene node. Many gene pair (but no mRNA isoform pair) are present in at least two tissue level gene non-functional networks.

**Table 7.** Summary statistics for single tissue gene level non-functional networks.

| Tissue | Genes | Gene pairs (Edges) | Tissue specific gene pairs | Density | Clusters | Largest connected component size | Gene pairs shared with other tissues |
| --- | --- | --- | --- | --- | --- | --- | --- |
| Adrenal glands | **(Nodes)** | 172220 | 167593 | 0.001514 | 218 | 14489 | 4627 |
| Adrenal glands | 14882 | 172220 | 167593 | 1.51E-03 | 218 | 14489 | 4627 |
| Embryonic facial prominence | 9601 | 8792 | 8464 | 1.84E-04 | 1438 | 5699 | 328 |
| Forebrain | 18556 | 1035497 | 1003429 | 5.83E-03 | 22 | 18520 | 32068 |
| Heart | 14025 | 26063 | 24758 | 2.52E-04 | 369 | 13342 | 1305 |
| Hindbrain | 18511 | 416933 | 396527 | 2.32E-03 | 36 | 18449 | 20406 |
| Kidney | 10024 | 10240 | 9678 | 1.93E-04 | 1234 | 7047 | 562 |
| Large intestine | 16997 | 87150 | 84028 | 5.82E-04 | 78 | 16880 | 3122 |
| Limb | 16981 | 100402 | 96023 | 6.66E-04 | 75 | 16877 | 4379 |
| Liver | 16600 | 102783 | 99668 | 7.23E-04 | 113 | 16427 | 3115 |
| Lung | 9358 | 8316 | 7991 | 1.83E-04 | 1562 | 4873 | 325 |
| Midbrain | 12586 | 15341 | 14363 | 1.81E-04 | 847 | 10728 | 978 |
| Neural tube | 10794 | 11410 | 10838 | 1.86E-04 | 1108 | 8191 | 572 |
| Ovary | 18573 | 715062 | 703048 | 4.08E-03 | 50 | 18497 | 12014 |
| Small intestine | 15084 | 84532 | 78142 | 6.87E-04 | 165 | 14833 | 6390 |
| Spleen | 12155 | 35158 | 33764 | 4.57E-04 | 559 | 11014 | 1394 |
| Stomach | 12764 | 23213 | 22020 | 2.70E-04 | 489 | 11829 | 1193 |

### Different mRNA isoforms of the same gene are functional in different tissues and have tissue preferred functional partners

The tissue level functional mRNA isoform networks along with the identification of gene pairs that are shared across tissues provide us an opportunity to distinguish the tissue-specific functional mRNA isoforms of a gene. We have identified around 164,000 functional gene pairs with different mRNA isoform pairs that are shared by multiple tissues. This points to the tissue specific expression and function of different mRNA isoforms of a gene.

The fraction of gene pairs shared between tissues is presented in Fig 9. We see that several pairs of tissues such as limb and forebrain, heart and large intestine, midbrain and forebrain, thymus and ovary, spleen and ovary etc. share a large number of gene pairs. This suggests that while these gene pairs are functional in multiple tissues, the actual mRNA isoform pairs can differ and our networks are capable of identifying such differential relationships between mRNA isoform pairs of the same gene pair.

The gene pair Fundc2 (FUN14 domain containing 2) and Necab1 (N-terminal EF-hand calcium binding protein 1) is present in both ovary and heart. The Fundc2 gene produces a single mRNA isoform NM_026126.4 while Necab1 gene produces two mRNA isoforms, XM_006538234.1 and NM_178617.4. The interaction between Fundc2 and Necab1 can be dissected into two interactions corresponding to the two mRNA isoform pairs (Fig 12A). Among the two mRNA isoform pairs, the pair involving XM_006538234.1 is heart specific functional mRNA isoform pair while the other pair involving NM_178617.4 is functional in ovary. This reveals the tissue preferred interaction partners of Fundc2 mRNA isoform NM_026126.4. Further investigation of all tissue specific functional mRNA isoform pairs involving Necab1 mRNA isoform XM_006538234.1 revealed that most of its interactions are found in heart (366 out of 391). Similarly, most of the interactions involving Necab1 mRNA isoform NM_178617.4 are found in ovary (836 out of 859). This highlights the expression and functional preference of Necab1 mRNA isoforms.

**Fig 12.**
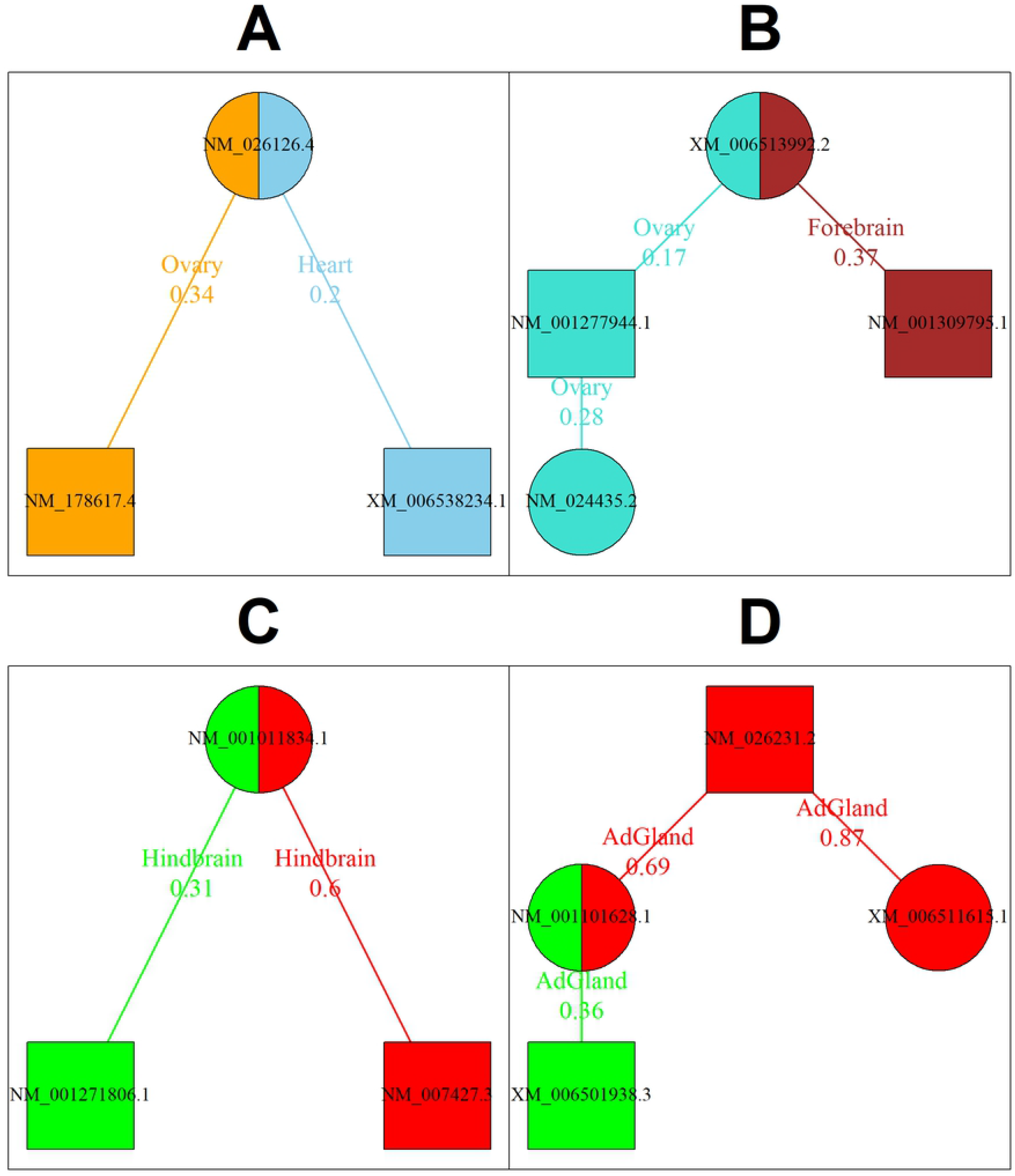
mRNA isoforms of the same gene have different functional partners across tissues. Few examples where the mRNA isoforms of the same gene have different functional/nonfunctional partners in specific tissues. The mRNA isoforms of the same gene are represented in same shape. The node color, edge color and the edge label color are encoded based on the tissue for part A and B. Functional pairs have green, while non-functional pairs have red node color, edge color and edge label color in parts C and D. Lower edge weight reflects higher strength of functional mRNA isoform pair. **A.** The mRNA isoform NM_026126.4 of gene Fundc2 forms a functional pair with different mRNA isoforms of Necab1 gene in heart and ovary. **B.** The ovary enriched mRNA isoform NM_001277944.1 of gene Apoc2 forms a functional pair with another ovary enriched Nts mRNA isoform NM_024435.2 in ovary. Other Apoc2 mRNA isoform NM_001309795.1 is preferred in forebrain. **C.** The Olfr1152 mRNA isoform NM_001011834.1 forms a functional pair with Agrp mRNA isoform NM_001271806.1 in hindbrain while the other pair involving Agrp mRNA isoform NM_007427.3.1 is non-functional in hindbrain. **D.** The gene pair Iqcf6 and Gstcd result in four mRNA isoform pairs of which one pair is functional and two are non-functional in adrenal glands.

Another such gene pair involves two mRNA isoform producing genes, Apoc2 (apolipoprotein CII) and Nts (neurotensin). The gene pair involving Apoc2 and Nts is found in the networks of ovary and forebrain and can be dissected into four interactions corresponding to the four mRNA isoform pairs. Three of these mRNA isoform interactions are found to be tissue-specific functional mRNA isoform pairs (Fig 12B). Interactions involving the Apoc2 mRNA isoform NM_001309795.1 are preferred in forebrain (1310 out of 1903) and NM_001277944.1 are preferred in ovary (355 out of 586). The NM_024435.2 mRNA isoform of Nts is enriched in ovary (1314 out of 1358) and interacts with the ovary enriched Apoc2 mRNA isoform NM_001277944.1 in ovary, suggesting a tissue preferred interaction pattern.

TENSION is also able to distinguish the tissue-specificity of mRNA isoforms of a gene between closely related tissues. For example, the gene Olfr994 (olfactory receptor 994) produces two mRNA isoforms, XM_006499549.1 and NM_146433.1. The mRNA isoform NM_146433.1 is preferred in hindbrain (223 out of 309 interactions) while XM_006499549.1 is preferred in midbrain (57 out of 65 interactions). There are several cases in which the mRNA isoforms of the same gene exhibit tissue preferred interactions. However, this is not true for all multi-isoform genes. The mRNA isoforms of many multi-isoform genes are not involved in tissue preferred interactions.

### Some mRNA isoform pairs are functional while other mRNA isoform pairs of the same gene pair are non-functional

We find about 660,000 instances where an mRNA isoform pair is functional while other mRNA isoform pairs of the same gene pair are non-functional. Around 143,000 of such cases are within the same tissue. For example, the mRNA isoforms of genes Agrp (agouti related neuropeptide) and Olfr1152 (olfactory receptor 1152) result in two mRNA isoform pairs (Fig 12C). The pair involving NM_001011834.1 (Olfr1152) and NM_001271806.1 (Agrp) is predicted to be functional in hindbrain while the other pair involving Agrp mRNA isoform NM_007427.3.1 is non-functional in hindbrain (Fig 12C). The NM_007427.3.1 mRNA isoform of Agrp is functionally enriched in the forebrain but has most of its non-functional interactions in hindbrain (362/447 functional interactions in forebrain vs 324/343 non-functional interactions in hindbrain), but the opposite is true for the isoform NM_001271806.1. The NM_001271806.1 mRNA isoform of Agrp contains an alternate 5’ exon, although both Agrp mRNA isoforms produce the same protein.

Similarly, for the gene pair involving Iqcf6 (IQ motif containing F6) and Gstcd (glutathione S-transferase, C-terminal domain containing), only one mRNA isoform pair is functional in adrenal glands while two other pairs are non-functional (Fig 12D). The remaining mRNA isoform pair could be functional or non-functional in multiple tissues.

The remaining 520,000 instances are across tissues, i.e., one mRNA isoform pair is tissue-specific functional in one tissue while other mRNA isoform pairs of the same gene pair are tissue-specific non-functional in other tissue.

### Validation of super-conserved and transcriptomic BodyMap of mouse tissue-specific genes

The first gene set contains 20 genes that are known to be widely expressed [50]. These genes have tissue-specific functional interactions in most of our 17 tissue-specific networks validating their ubiquitous expression and function (Fig. 13). The second gene set contains 1654 genes from the transcriptomic BodyMap of mouse that have a very high expression in one tissue (relative to all other tissues) and thus a higher propensity to have more tissue-specific functions. For every gene, we compute the top *n* = {1, 3, 5, 7, 9, *All*} tissues for its mRNA isoforms based on the number of functional interactions in the tissue.

**Fig 13.**
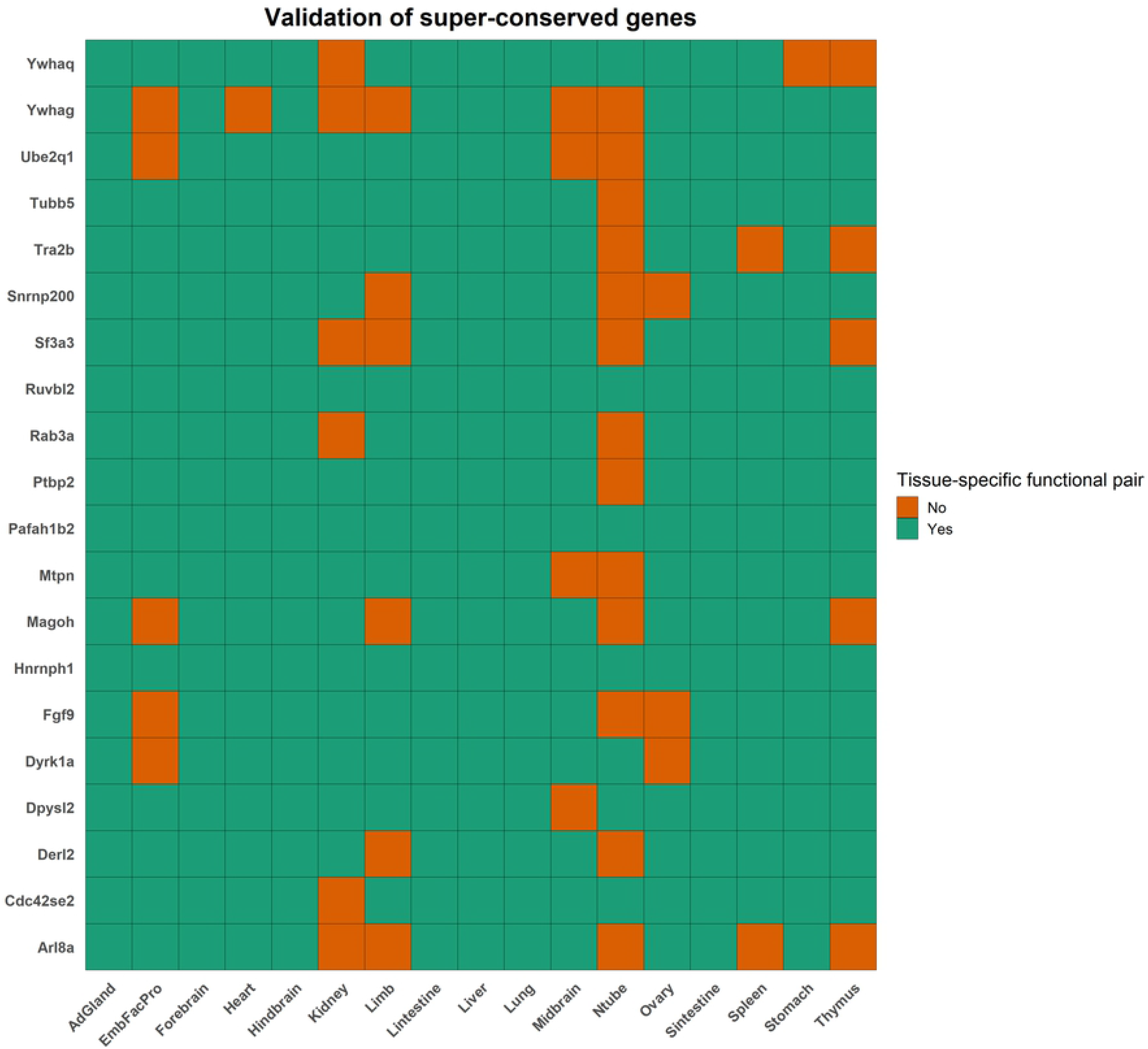
Validation of super-conserved genes. A heatmap showing the presence or absence of a tissue-specific functional interaction for the 20 super-conserved genes. The genes are on the y-axis and the tissues are on the x-axis. If a gene has a tissue-specific functional interaction, the corresponding block is filled green, or orange otherwise. Abbreviations - AdGland: Adrenal glands; EmbFacPro: Embryonic Facial Prominence; Lintestine: Large intestine; Ntube: Neural tube; Sintestine: Small intestine.

We find that the top tissue (*n* = 1) among our tissue-specific networks and that in the transcriptomic BodyMap of mouse matches for 503 genes (30%; Table 7). However, a gene can be involved in multiple functions across multiple tissues due to different mRNA isoforms. Therefore, when we consider the top 3 (52% match) or top 5 (68% match) tissues, we find a much higher correlation with the transcriptomic BodyMap of mouse (Table 7). Overall, we find 1245 (75%) genes to have at least one tissue specific interaction in the same tissue as described in the transcriptomic BodyMap of mouse.

It is interesting to note that if we consider the tissue-specificity of only the genes, ignoring the tissue-specificity of different mRNA isoforms of the same gene, we find a weaker correlation with the transcriptomic BodyMap of mouse (15% and 41% respectively for n = 1 and 3). Most studies including the transcriptomic BodyMap of mouse focus only on the gene expression and function, completely ignoring the effects of alternatively spliced mRNAs. Our study further illustrates the importance of distinguishing the functions of different mRNA isoforms of the same gene.

**Table 8.**
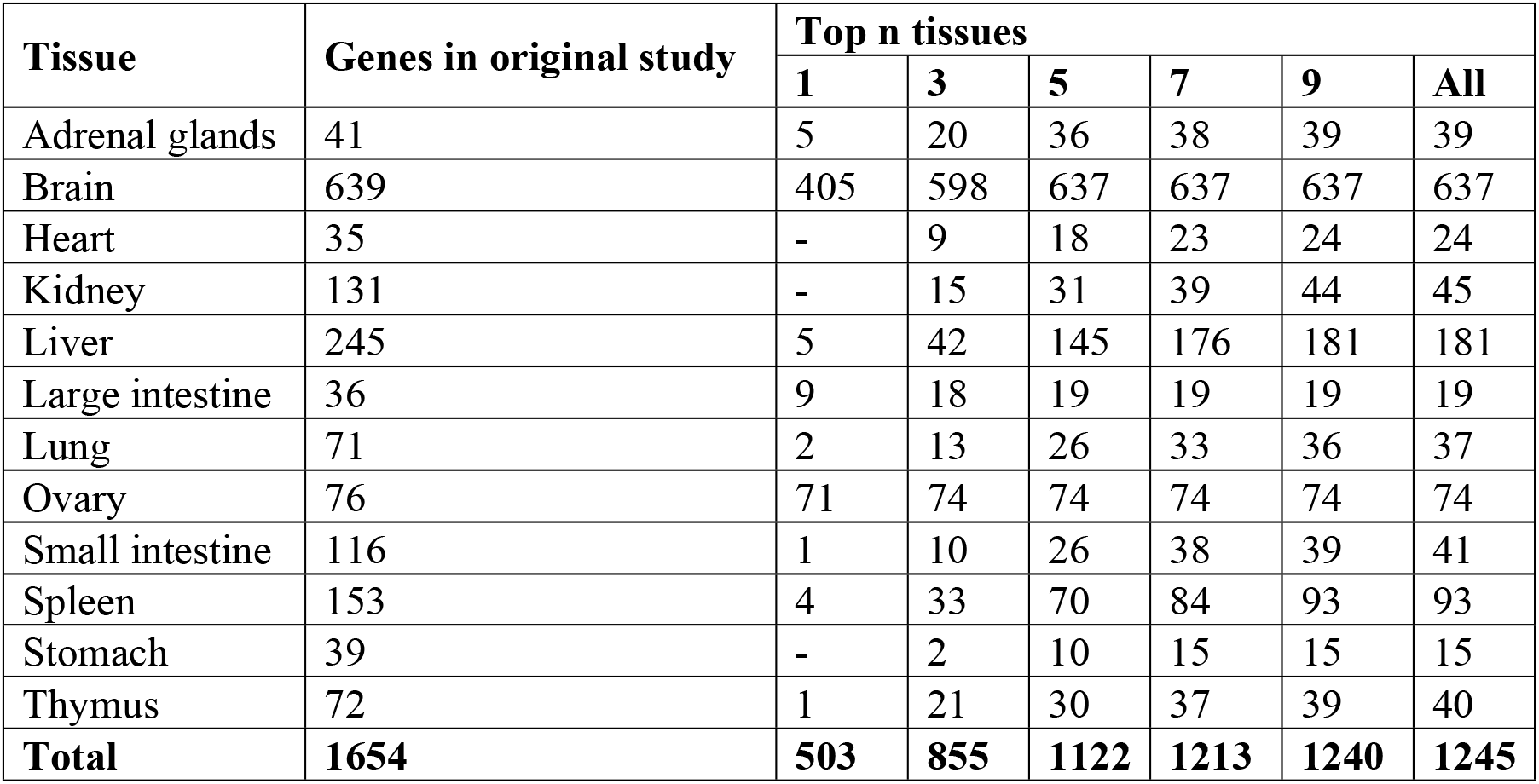
Number of genes with tissue-specific functional interactions in the same tissue as the transcriptomic BodyMap of mouse.

| Tissue | Genes in original study | Top n tissues |  |  |  |  |  |
| --- | --- | --- | --- | --- | --- | --- | --- |
|  |  | 1 | 3 | 5 | 7 | 9 | All |
| Adrenal glands | 41 | 5 | 20 | 36 | 38 | 39 | 39 |
| Brain | 639 | 405 | 598 | 637 | 637 | 637 | 637 |
| Heart | 35 | - | 9 | 18 | 23 | 24 | 24 |
| Kidney | 131 | - | 15 | 31 | 39 | 44 | 45 |
| Liver | 245 | 5 | 42 | 145 | 176 | 181 | 181 |
| Large intestine | 36 | 9 | 18 | 19 | 19 | 19 | 19 |
| Lung | 71 | 2 | 13 | 26 | 33 | 36 | 37 |
| Ovary | 76 | 71 | 74 | 74 | 74 | 74 | 74 |
| Small intestine | 116 | 1 | 10 | 26 | 38 | 39 | 41 |
| Spleen | 153 | 4 | 33 | 70 | 84 | 93 | 93 |
| Stomach | 39 | - | 2 | 10 | 15 | 15 | 15 |
| Thymus | 72 | 1 | 21 | 30 | 37 | 39 | 40 |
| **Total** | **1654** | **503** | **855** | **1122** | **1213** | **1240** | **1245** |

### Similar tissues have similar mRNA isoform expression profile

Tissues that are functionally and morphologically similar tend to have more consistent gene expression profile than other tissues [51]. We also observe that similar tissues such as midbrain, forebrain, hindbrain and neural tube have a very high Pearson correlation coefficient (ρ ≥ 0.97; Fig 13) based on the median mRNA isoform expression profile. Likewise, adrenal gland is most highly correlated with ovary (ρ = 0.87), large intestine with small intestine (ρ = 0.84) and thymus with spleen (ρ = 0.88) among others, and are consistent with previous findings [51].

**Fig 14.**
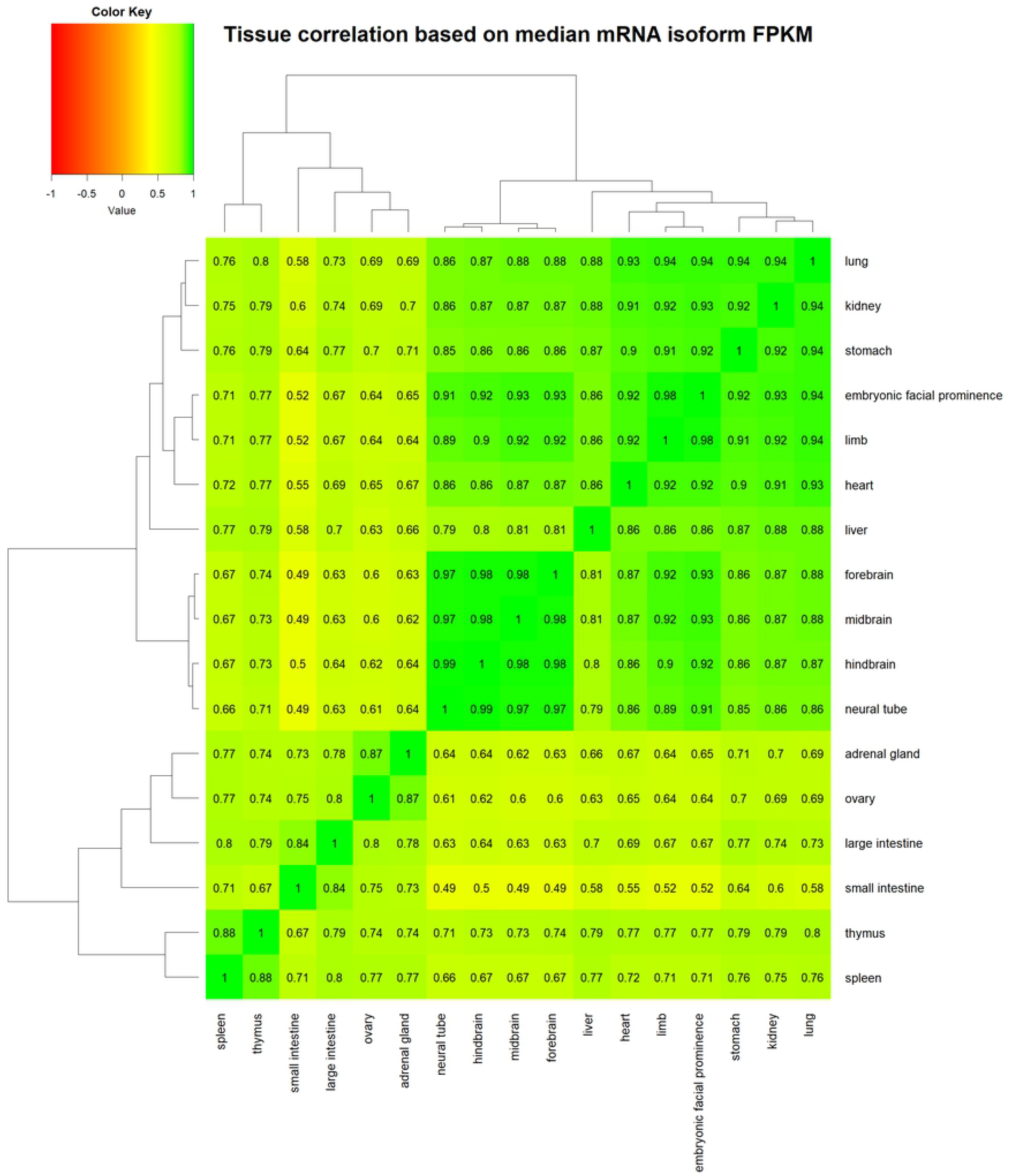
Similar tissues have similar mRNA isoform expression profile. A heatmap showing the Pearson correlation coefficient between pairs of tissue based on the median mRNA isoform expression values. The dendrogram on the rows and columns reflects the clustering of tissues. Green represents higher positive correlation between a pair of tissue while red reflects higher negative correlation. Similar tissues can be seen being clustered together.

## Discussions

We have developed tissue-level functional networks to study mRNA isoform functional relationships, providing a higher resolution view of biological processes as compared to traditional gene-level networks. Learning the differences in the functional connections of mRNA isoforms of the same gene are crucial for functional genomics, and helps us in deepening our understanding of gene functions. Determining the functional interaction patterns of mRNA-isoforms of the same gene also provides useful information about biological regulation, diseases, and stress response caused by AS.

It is widely believed that the fate of biological processes and pathways varies with different mRNA isoforms of the same gene. Many pathways and molecular processes differ across cell and tissue-types. These mechanisms are also altered by external conditions such as abiotic and biotic stress. Understanding of such deviations in cell, tissue and condition specific functional relationships would be of interest to understand the perturbed mechanisms.

Based on the analysis of 359 mouse tissue-specific RNA-Seq samples along with 9 diverse sequence properties, we have constructed 17 tissue-specific mRNA isoform level functional networks. These networks constitute ~10.6 million unique functional and ~3.5 million nonfunctional mRNA isoform interactions across 17 tissues. In addition to these tissue-specific networks, we have also developed an organism-wide reference network. We show that TENSION is highly accurate with very high precision and recall by comparing our predictions with class label shuffled datasets, ten-fold stratified cross validation, previous method, and updated annotations from gene ontology, pathway databases and PPIs. In addition to these, we also validate our predictions by using a gene set of 20 ubiquitously expressed genes and 1654 genes with a very high expression in one tissue from the transcriptomic BodyMap of mouse. The improvement in the performance (compared to the original study) of Bayesian network based MIL method on our dataset also prove the utility of TENSION in generating better mRNA isoform level datasets.

Our tissue-specific networks capture the differences in functional relationships of mRNA isoforms of the same gene across multiple tissues highlighting the importance of tissue-specific changes in biological processes and pathways. We are also able to distinguish the tissue-specific functional mRNA isoforms of a gene. We also find that different mRNA isoforms of the same gene are enriched in different tissues, suggesting differential tissue-level activity of mRNA isoforms of the same gene. Furthermore, we also see that morphologically and functionally similar tissues tend to have more consistent mRNA isoform expression profile.

By studying the gene level networks in conjunction with mRNA-isoform level functional networks, we are able to gain different insights into the molecular mechanisms of biological processes. Diving down further into the tissue-specific networks sheds more light on the tissue-level activities of a gene and its mRNA isoforms. The central genes identified in these tissue-level networks are enriched in tissue related processes.

In summary, we provide the research community with a comprehensive characterization of mRNA isoform level tissue-specific functional networks for mouse. TENSION is simple and generic, making it easily applicable to other organisms. We expect that these networks will allow further in-depth investigations of the impact of alternatively spliced mRNA isoforms on biological processes. We anticipate that tissue-specific mRNA-isoform functional networks will find wide applications in genomics, agriculture and biomedical sciences.

## Data availability

All data and scripts will be available at DataShare: Iowa State University’s Open Research Data Repository.

## Acknowledgement

This work used the Extreme Science and Engineering Discovery Environment (XSEDE), which is supported by National Science Foundation grant number ACI-1548562. This work used the XSEDE Comet cluster at San Diego Supercomputer Center (SDSC) through allocation TG-BIO170049. We would like to thank Dr. Mahidhar Tatineni and Dr. Martin Kandes from SDSC User Services group for providing technical support at Comet cluster. We also thank Megan O’Donnell and Levi Baber for setting up the data repository at DataShare: Iowa State University’s Open Research Data Repository. We would also like to thank Dr. Yuanfang Guan and Dr. Hongdong Li for providing some data and the code for the comparison with Bayesian network based MIL method.

---

**Fig S1.**
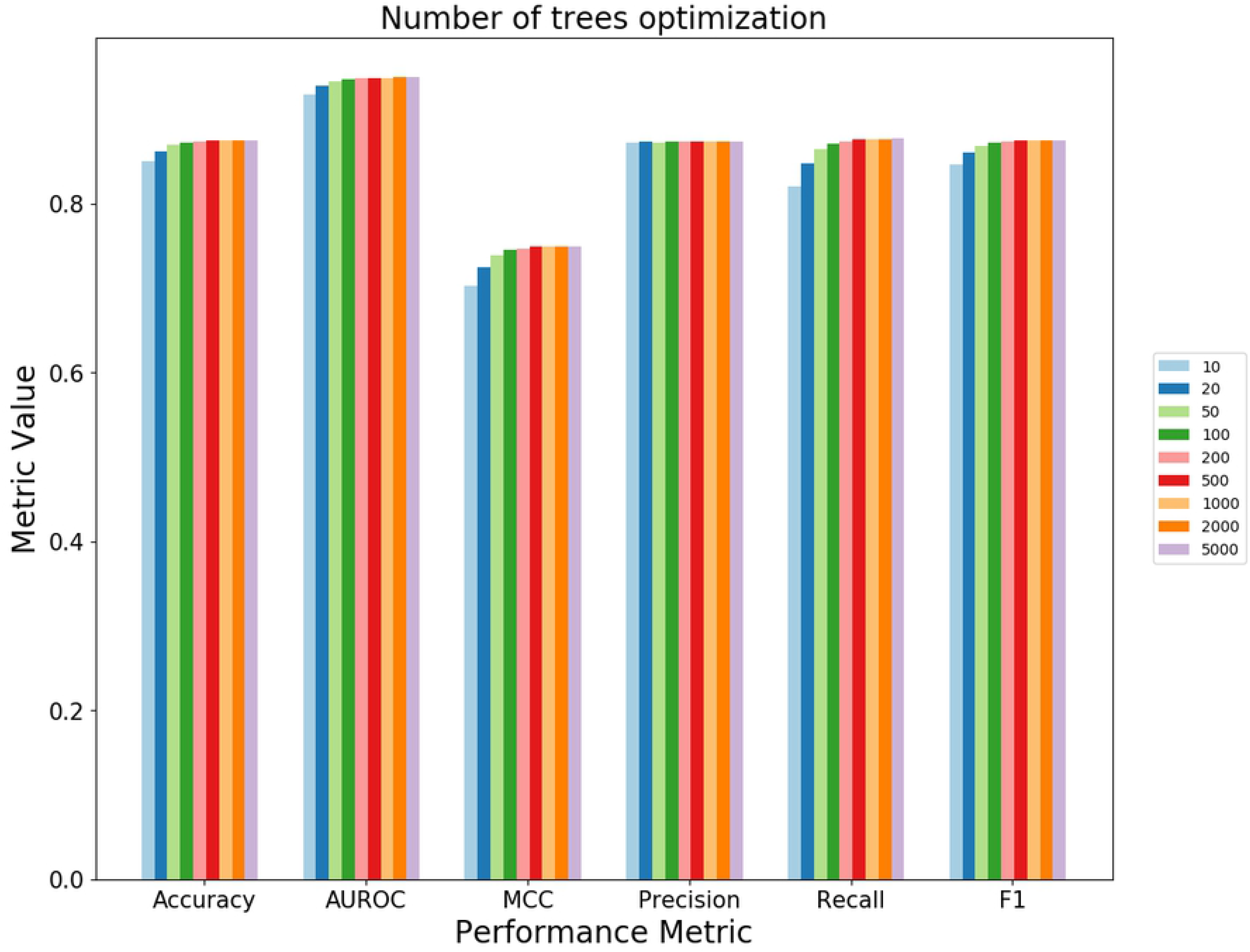
Optimization of number of trees in random forest. A bar plot of performance metrics computed at different number of trees used in random forest. There is very little improvement in the performance beyond 100 trees, therefore, we have used 100 trees while developing TENSION. Abbreviations - ROC: Receiver Operating Characteristic; MCC: Matthews Correlation Coefficient.

**Fig S2.**
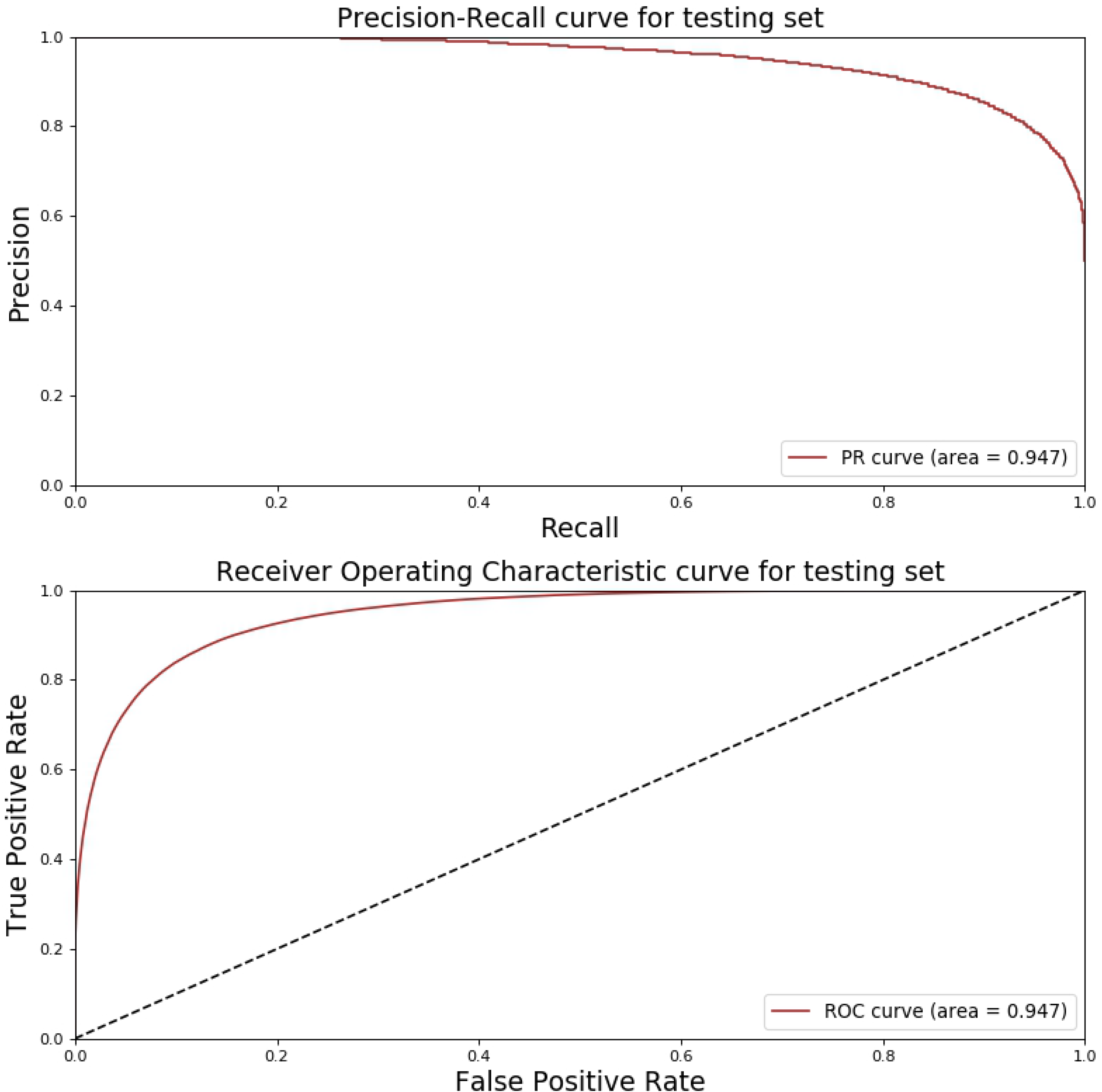
Performance evaluation on original testing dataset. The precision-recall and receiver operating characteristic curve for predictions on the original testing dataset. A model with area under the curve closer to 1 is better while a model with an area under the curve of 0.5 is equivalent to making random guess. Abbreviations - PR: Precision-Recall; ROC: Receiver Operating Characteristic.

**Table S1. List of RNA-Seq experiments and their tissue.**

